# Makorin 1 controls embryonic patterning by alleviating Bruno1-mediated repression of *oskar* translation

**DOI:** 10.1101/501643

**Authors:** Annabelle Dold, Hong Han, Niankun Liu, Andrea Hildebrandt, Mirko Brüggemann, Cornelia Rücklé, Anke Busch, Petra Beli, Kathi Zarnack, Julian König, Jean-Yves Roignant, Paul Lasko

**Affiliations:** RNA Epigenetics, Institute of Molecular Biology, 55128 Mainz, Germany; Department of Biology, McGill University, Montréal, Québec, Canada H3G 0B1; Chromatin Biology and Epigenetics, Institute of Molecular Biology, 55128 Mainz, Germany; Genomic Views of Splicing Regulation, Institute of Molecular Biology, 55238 Mainz, Germany; Buchmann Institute for Molecular Life Sciences, Frankfurt, Germany; Bioinformatics Core Facility, Institute of Molecular Biology, 55128 Mainz, Germany

**Keywords:** Mkrn1, Bruno, oskar, polyadenylation, pAbp, oogenesis, germ cell specification, pole plasm, *Drosophila*

## Abstract

Makorins are evolutionary conserved proteins that contain C_3_H-type zinc finger modules and a RING E3 ubiquitin ligase domain. In *Drosophila* maternal Makorin 1 (Mkrn1) has been linked to embryonic patterning but the mechanism remained unsolved. Here, we show that Mkrn1 is essential for axis specification and pole plasm assembly by translational activation of *oskar*. We demonstrate that Mkrn1 interacts with poly(A) binding protein (pAbp) and binds *osk* 3’ UTR in a region adjacent to A-rich sequences. This binding site overlaps with Bruno1 (Bru1) responsive elements (BREs), which regulate *osk* translation. We observe increased association of the translational repressor Bru1 with *osk* mRNA upon depletion of Mkrn1, indicating that both proteins compete for *osk* binding. Consistently, reducing Bru1 dosage partially rescues viability and Osk protein level in ovaries from *Mkrn1* females. We conclude that Mkrn1 controls embryonic patterning and germ cell formation by specifically activating *osk* translation by displacing Bru1 from its 3’ UTR.

**Author Summary:** To ensure accurate development of the *Drosophila* embryo, proteins and mRNAs are positioned at specific sites within the embryo. Many of these proteins and mRNAs are produced and localized during the development of the egg in the mother. One protein essential for this process that has been heavily studied is Oskar (Osk), which is positioned at the posterior pole. During the localization of *osk* mRNA, its translation is repressed by the RNA-binding protein Bruno1 (Bru1), ensuring that Osk protein is not present outside of the posterior where it is harmful. At the posterior pole, *osk* mRNA is activated through mechanisms that are not yet understood. In this work, we show that the conserved protein Makorin 1 (Mkrn1) is a novel factor involved in the translational activation of *osk*. Mkrn1 binds specifically to *osk* mRNA in a region that overlaps with the binding site of Bru1, thus alleviating the association of Bru1 with *osk*. Moreover, Mkrn1 is stabilized by poly(A) binding protein, a translational activator that binds *osk* mRNA in close proximity to Mkrn1. Our work thus helps to answer a long-standing question in the field, providing insight about the function of Mkrn1 and more generally into embryonic patterning in animals.

## Introduction

In the *Drosophila* embryo, the maternally deposited pole plasm is a site of specialized translation of mRNAs required for germ cell specification and posterior patterning [1]. Numerous mRNAs accumulate in the pole plasm during oogenesis and early embryogenesis through several different localization mechanisms [2,3]. Among these mRNAs is *oskar* (*osk*), which localizes during oogenesis to the posterior along a polarized microtubule network [4] and via a trapping mechanism [5]. Several lines of evidence indicate that *osk* is the primary determinant that specifies germ cells and posterior patterning. Ectopic expression of *osk* at the anterior can induce a second set of pole cells and a bicaudal embryonic segmentation pattern with mirror-image posterior segments [6,7]. Mutations such as *Bicaudal-D* (*Bic-D*), *ik2*, and others that produce a duplicated anterior focus of *osk* mRNA also produce bicaudal embryos [8-10]. Conversely, embryos from females carrying hypomorphic loss-of-function mutations of *osk* lack posterior segmentation and pole cells [11]. Mutations in a number of other genes can produce a similar phenotype, and these are collectively known as posterior-group genes [12]. Some of these genes (for example *cappuccino, chickadee, spire and staufen* (*stau*)) are required for posterior localization of *osk*. A failure to deploy *osk* produces the posterior-group phenotype in these mutants [13,14]. Other posterior-group genes (for example *vasa* (*vas*), *tudor, nanos* (*nos*), and *aubergine* (*aub*)), produce mRNAs and/or proteins that also accumulate in pole plasm and operate downstream of *osk* [15-17].

*osk* translation is under elaborate temporal and spatial regulation, ensuring that Osk protein becomes abundant only in the posterior pole plasm and not before stage 9 of oogenesis [18]. A key repressor of *osk* translation prior to that stage and outside the pole plasm is Bruno1 (Bru1), which interacts with binding sites called Bru1 response elements (BREs) in the *osk* 3’ UTR. Mutations affecting the BREs result in premature and excessive Osk expression [18,19]. There is evidence for two distinct mechanisms of translational repression of *osk*. In the first model, Bru1 recruits Cup, which inhibits assembly of an active cap-binding complex by competitively inhibiting eIF4G for binding to eIF4E [20]. An alternative model involves oligomerization of *osk* mRNA into large ribonucleoprotein particles (RNPs) that are inaccessible to the translational machinery [21-23]. These mechanisms may be connected, in that physically concentrating *osk* mRNA molecules into RNPs can enable regulation in *trans* through inter-molecular interactions. For instance, Bru1 bound to the 3’ end of one *osk* mRNA molecule could recruit Cup to eIF4E bound to the 5’ cap structure of another *osk* mRNA molecule, thus repressing its translation [24,25]. Importantly, *osk* translation and *osk* mRNA localization are tightly coupled. In nonsense *osk* alleles or when 3’ UTR elements required for translational activation are mutated, *osk* mRNA localizes transiently to the pole plasm but its accumulation is not maintained [9,26,27]

While several proteins have been implicated in activating *osk* translation in the pole plasm [14,28,29], a comprehensive picture of how this is achieved has not yet emerged. For instance, it has been proposed that activation of *osk* translation involves inhibition of Bru1 [23]. Related to this, a BRE-containing region in the distal part of the *osk* 3’ UTR (BRE-C) functions in repression as well as in activation [24]. Nevertheless, the mechanism underlying the dual function of this element has not yet been solved.

Large-scale *in situ* hybridization screens have identified many other mRNAs that localize to the pole plasm [3,30], and some of the corresponding genes could potentially also be involved in *osk* regulation. To search for new posterior-group genes, we previously expressed shRNAs targeting 51 different mRNAs that accumulate in the pole plasm to determine if doing so produced defects in posterior patterning or pole cell formation. We observed that a substantial proportion of embryos produced by *Makorin 1* (*Mkrn1*) knockdown females showed a posterior-group phenotype [31]. Makorin proteins are conserved in plants, fungi, and animals, and contain a RING-domain as well as one or more C_3_H-type zinc fingers (ZnF) [32]. The role of makorin proteins is somewhat enigmatic despite their widespread evolutionary conservation. Mammalian MKRN1 has been identified as a E3 ubiquitin ligase that promotes degradation of target proteins [33], but proteomic analysis does not support an association with proteasome components [34]. Furthermore MKRN1’s shorter isoform stimulates translation in rat forebrain neurons but the mechanism is currently unknown [35,36].

Here, we analyzed the function of Mkrn1 during oogenesis and early embryogenesis. We generated several alleles that alter different domains of the *Mkrn1* coding sequence. Using these mutants, we found that Mkrn1 is required for accumulation of Osk protein at the posterior pole of the oocyte. Furthermore, we present evidence that Mkrn1 directly binds to a site in the *osk* 3’ UTR that overlaps the BRE-C domain via its N-terminal zinc finger domain. This binding site is adjacent to an A-rich region that recruits pAbp to the 3’ UTR [37]. The association between Mkrn1 and *osk* mRNA is stabilized by physical association with pAbp. Strikingly, depletion of *Mkrn1* results in an increased level of Bru1 binding to *osk* mRNA, and reduction of *bru1* gene dosage partially rescues *Mkrn1* mutant phenotypes. Based on this evidence we propose that Mkrn1 competes with Bru1 for *osk* mRNA binding, thus positively regulating *osk* translation and explaining the specific role of BRE-C in translational activation.

## Results

### The *Drosophila* genome includes four Makorin-related genes

In many organisms, up to four distinct genes encoding members of the Makorin family exist, but only one such gene, *Mkrn1*, has been annotated in *Drosophila*. To investigate whether flies are unusual in this regard, we searched for sequences similar to human MKRN1. This analysis uncovered four *Drosophila* genes, namely *Mkrn1, CG5334, CG5347*, and *CG12477*, with substantial similarity to *MKRN1* (Fig S1A). All four predicted polypeptides from these genes contain a region of approximately 130 amino acids that is highly conserved and contains a RING-domain as well as C_3_H-type zinc fingers (ZnF). The proteins are otherwise more divergent from one another, with the exception that all but CG12477 contain a ZnF domain near the amino-terminus.

To analyze the differences in their functionalities, we first determined the expression profile of all four *Makorin* genes during development. *Mkrn1* mRNA is expressed at detectable levels at all developmental stages (Fig 1A) and clearly peaks in early (0 - 2.5 h) embryos and ovaries. In contrast, expression of the other three Makorin genes is undetectable during early development but peaks in adult males (Fig S1B). Together, these results indicate that *Mkrn1* is the only gene of the family expressed in ovaries and early embryos and suggest that the three other genes could be specifically expressed in testes.

**Figure 1.**
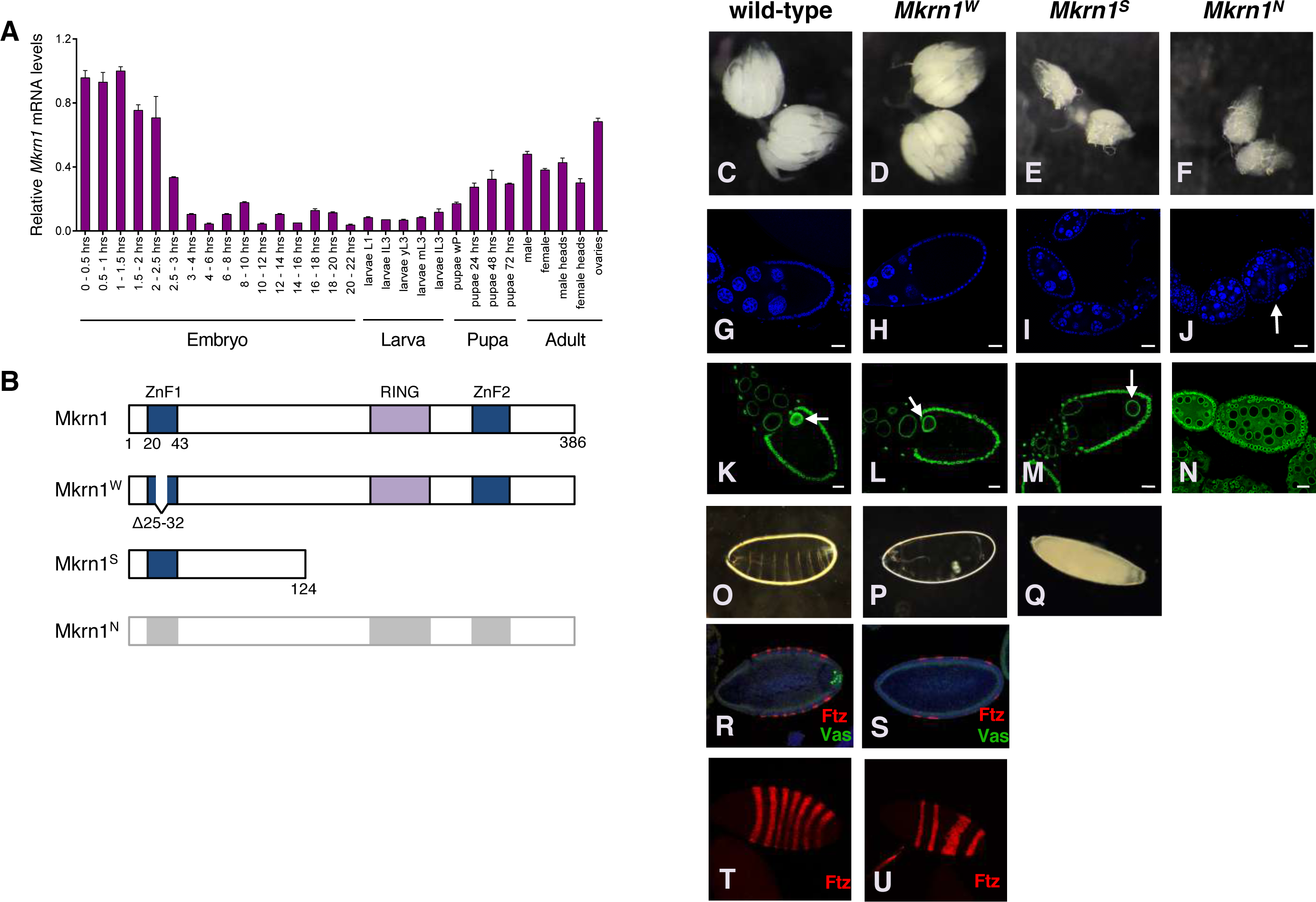
*Mkrn1* alteration affects ovarian development. Relative *Mkrn1* mRNA levels (normalized to *Rpl15* mRNA) at various stages of development, as measured by quantitative RT-PCR. Error bars depict Stdev, n=3. (**B**) Schematic diagram of the proteins encoded by the *Mkrn1* alleles used to analyze its function in vivo. *Mkrn1*^*N*^ is a null allele and produces no protein. (**C**-**F**) Bright-field micrographs of entire ovaries from wild-type and *Mkrn1* mutants. Note the reduced size of *Mkrn1*^*S*^ and *Mkrn1*^*N*^ ovaries. (**G**-**J**) Individual egg chambers stained with the DNA marker DAPI. Fewer stage-10 and older egg chambers are present in *Mkrn1*^*S*^ and no late stage egg chambers are present in *Mkrn1*^*N*^ ovaries. Abscission defects resulting from inappropriate follicle cell migration are frequently observed in *Mkrn1*^*N*^ ovaries (**J**, arrow). (**K**-**N**) Individual egg chambers stained with α-Lamin to highlight nuclear membranes. (**M**) The oocyte nucleus (marked with an arrow in **K**, **L**, and **M**) remains at the posterior of *Mkrn1*^*S*^ oocytes. (**N**) Some *Mkrn1*^*N*^ egg chambers have 16 germline cells whose nuclei are all of similar size, suggesting a defect in oocyte differentiation. Note also irregularities in the follicle cell monolayer in the *Mkrn1*^*N*^ egg chamber. (**O**-**Q**) Dark field photographs of eggs and embryos produced by wild-type and *Mkrn1* mutants. (**P**) Most embryos produced by *Mkrn1*^*W*^ females have a posterior-group phenotype. (**Q**) Eggs produced by *Mkrn1*^*S*^ females lack dorsal appendages and do not support embryonic development. Scale bars, 20 μm. (**R**, **S**) Immunostaining with α-Ftz (red) and α-Vas (green) reveals segmentation defects and the absence of pole cells in *Mkrn1*^*W*^ embryos. (**T**, **U**) Immunostaining with α-Ftz (red) of wild-type and *Mkrn1*^*W*^ embryos imaged on a surface focal plane to better visualize striped patterns of expression.

### *Mkrn1* mutants reveal essential roles in oogenesis and embryogenesis

To elucidate the role of *Mkrn1*, we used CRISPR/Cas9 to produce three different mutant alleles: *Mkrn1*^*N*^, a complete deletion of the coding sequence, *Mkrn1*^*S*^, a frameshift mutation that is predicted to produce a C-terminally truncated protein of 124 amino acids, including only ZnF1 among conserved domains, and *Mkrn1*^*W*^, a small in-frame deletion that disrupts only the ZnF1 domain (Fig 1B). In the strong *Mkrn1* mutants (*Mkrn1*^*S*^ *and Mkrn1*^*N*^), most egg chambers cease development at or before stage 10 (Figs 1C-J). The nuclei of *Mkrn1*^*S*^ oocytes that progress as far as stage 9 or later remain at the posterior, failing to migrate to the antero-dorsal corner (Fig 1M). The very few eggs laid by *Mkrn1*^*S*^ females have no dorsal appendages and do not develop (Fig 1Q). *Mkrn1*^*N*^ egg chambers do not progress as far as stage 9 and showed variable defects in early oogenesis including failure of oocyte differentiation (Fig 1N) and inappropriate follicle cell migration (Figs 1J and N). On the other hand, ovaries of females homozygous for *Mkrn1*^*W*^ have a similar morphology to wild-type (Figs 1C and D). *Mkrn1*^*W*^ mutant ovaries completed oogenesis and produced fertilizable eggs in similar numbers as wild-type controls (Figs 1G, H, K and L, and Table 1). To examine the role of *Mkrn1* in embryonic patterning, we compared cuticle preparations of wild-type and *Mkrn1*^*W*^ embryos (Figs 1O, P). We found that most *Mkrn1*^*W*^ embryos lack posterior segments, a phenotype similar to *osk* mutants and to what we previously observed at lower frequency from females expressing shRNA targeting *Mkrn1* [31]. To investigate this more closely, we stained wild-type and *Mkrn1*^*W*^ embryos for Fushi tarazu (Ftz) and Vas proteins. In wild-type blastoderm-stage embryos, Ftz is expressed in seven stripes along the anterior-posterior axis, while in posterior-group embryos the number of Ftz stripes is usually reduced to four [38] (Fig 1R, 1T). At blastoderm stage in wild-type embryos, Vas-positive pole cells are clustered at the posterior pole [16] (Fig 1R). Consistent with a posterior-group phenotype, we observed four Ftz stripes in 65% of *Mkrn1*^*W*^ embryos (30/46) (Fig 1S, 1U), with most of the rest showing fewer Ftz stripes with an additional broad domain of Ftz expression. In addition, 95% of *Mkrn1*^*W*^ embryos (63/66) showed no pole cells by Vas staining (Fig 1S). The posterior patterning defects result in lethality for most (97%) *Mkrn1*^*W*^ embryos. However, the small number of *Mkrn1*^*W*^ embryos that hatch into viable larvae (40/1222, 3.3%, Table 1) can complete development to adulthood.

**Table 1.**
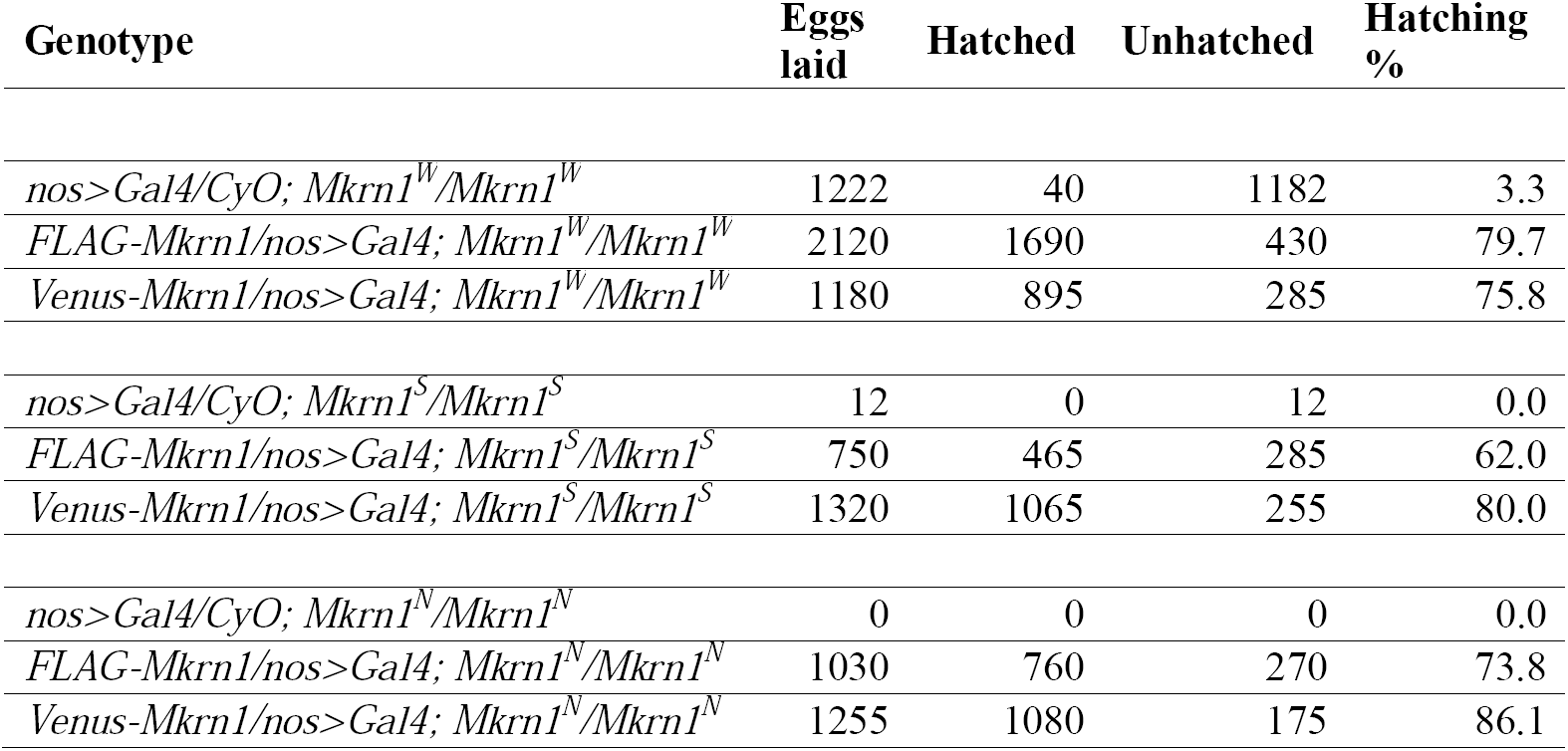
Expression of tagged *Mkrn1* from transgenes rescues oogenesis and viability to embryos produced by *Mkrn1* mutant females.

### Mkrn1 accumulates in pole plasm during oogenesis

To examine the distribution of Mkrn1 in the germline, we expressed transgenic Venus- and FLAG-tagged Mkrn1 using the *nos>GAL4* driver. Since *Mkrn1* females could be rescued to fertility by germline specific expression of these transgenes (Table 1), we concluded that these tagged transgenes are functional, and thus inferred that their localization should reflect the endogenous one. When expressed in ovaries, Venus- or FLAG-tagged Mkrn1 (subsequently called Mkrn1) becomes detectable in a uniform distribution in germline cells from early oogenesis. We observed a mild accumulation of Mkrn1 in cytoplasmic particles resembling nuage at the outer surface of nurse cell nuclear membranes and in the early oocyte (Fig 2A, Figs S2A and S). In later egg chambers Mkrn1 remains abundant in nurse cells but tightly localized in the pole plasm in the oocyte (Figure 2D). Next, we conducted double labelling experiments in wild-type ovaries to determine the degree of colocalization between Mkrn1 and known pole plasm components. In both, stage 8 and stage 10 oocytes, Mkrn1 co-localizes extensively with Stau (Figs 2A-F, Fig S2A-F), *osk* mRNA (Figs 2G-L), Osk protein (Figs S2G-L), Vas (Figs S2M-R) and Aub (Figs S2S-X). This close association between Mkrn1 and many important pole plasm components suggests that Mkrn1 is an integral component of pole plasm.

**Figure 2.**
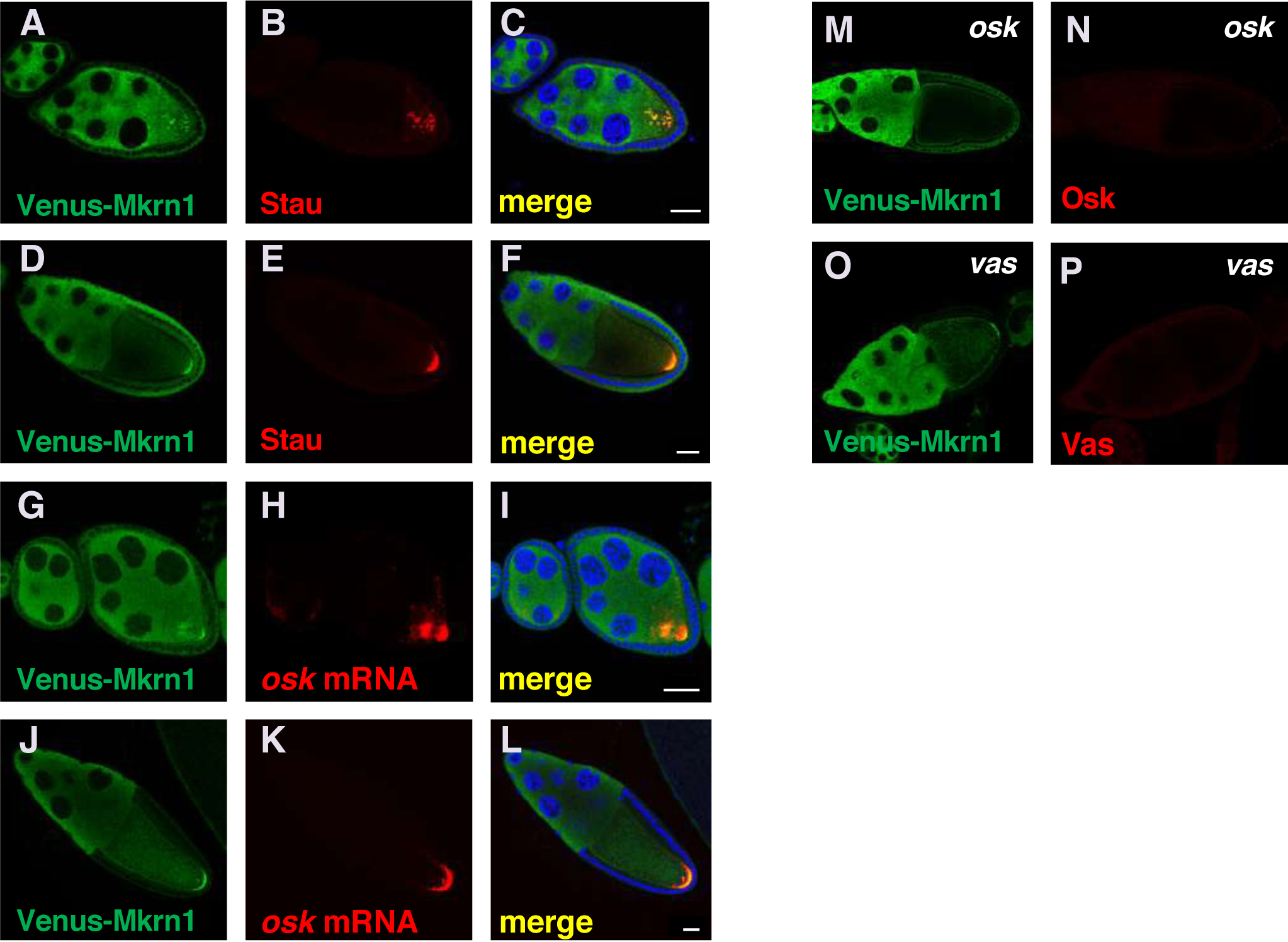
Mkrn1 accumulates in pole plasm. (**A**-**C**) The three panels show the same egg chambers stained for (**A**) Venus-Mkrn1, Stau, and a (**C**) merged image. Venus-Mkrn1 expression was driven by *nos*>Gal4. Colocalization of Venus-Mkrn1 and Stau can be observed in particles that have not yet accumulated at the posterior of the early stage 8 oocyte. (**D**-**F**) The three panels show the same stage 10 egg chamber stained for (**D**) Venus-Mkrn1, (**E**) Stau and (**F**) a merged image. There is extensive colocalization of Venus-Mkrn1 and Stau in the posterior pole plasm of the oocyte. (**G**-**I**) The three panels show the same egg chambers stained for (**G**) Venus-Mkrn1, (**H**) *osk* mRNA, and (**I**) a merged image. Colocalization of Venus-Mkrn1 and *osk* can be observed in an early stage 8 oocyte where *osk* has not yet fully localized at the posterior of the oocyte. (**J**-**L**) The three panels show the same stage 10 egg chamber stained for (**J**) Venus-Mkrn1, (**K**) *osk* mRNA and (**L**) a merged image. There is extensive colocalization of Venus-Mkrn1 and *osk* mRNA in the posterior pole plasm of the oocyte. (**M**, **N**) Venus-Mkrn1 expressed in an *osk*^*54*^*/Df(3R)p-XT103* genetic background. Venus-Mkrn1 fails to accumulate in pole plasm. (**O**, **P**) Venus-Mkrn1 expressed in a *vas*^*1*^*/vas*^*PH*^ genetic background. Venus-Mkrn1 accumulates normally in pole plasm. Scale bars, 25 μ m.

To determine whether Mkrn1 depends on the pole plasm assembly pathway for its posterior localization, we expressed the tagged *Mkrn1* transgenes in *osk* (*osk*^*54/Df*^) and *vas* (*vas*^*1*^/*vas*^*PH*^) mutant backgrounds. We found that loss of *osk* abolished Mkrn1 localization (Figs 2M-N, Fig S3A). In contrast, Mkrn1 localized normally to the posterior in *vas* mutant oocytes (Figs 2O-P, Fig S3B), placing Mkrn1 between *osk* and Vas in the pole plasm assembly pathway.

To obtain insights into the link between Mkrn1 and pole cell determination, we collected embryos from females trans-heterozygous for a *Mkrn1* allele and for either a *vas* or *osk* allele. Next, we compared the number of pole cells with single heterozygous controls. When heterozygous, *Mkrn1*^*W*^ or *Mkrn1*^*S*^ had little effect on pole cell number. However, either allele reduced the number of pole cells produced by *vas*^*PH*^ heterozygotes, and further by *osk*^*54*^ heterozygotes (Fig S4A). These data support a genetic interaction between *Mkrn1* and genes involved in embryonic patterning and pole cell specification.

### Mkrn1 ensures correct deployment of specific mRNAs and proteins involved in embryonic patterning

To address whether *Mkrn1*mutations may affect the distribution of proteins involved in embryonic patterning, we performed immunostaining experiments. We found a striking reduction in posterior accumulation of Osk in oocytes from all *Mkrn1* alleles (Figs 3A-D). For Stau, we observed weaker and more diffuse posterior localization in *Mkrn1*^*W*^ as compared to wild-type (Figs 3E-F, Figs S4B-D) and no localized protein in *Mkrn1*^*S*^ and *Mkrn1*^*N*^ (Figs 3G and H). On the other hand, Grk localized normally to the antero-dorsal corner of the oocyte in *Mkrn1*^*W*^ (Figs 3I-J, Figs S4E-G). In *Mkrn1*^*S*^, Grk was observed at reduced levels associated at the posterior with the mislocalized oocyte nucleus (Fig 3K, Figs S4H-J) and diffusely distributed at a very low level in *Mkrn1*^*N*^ (Fig 3L). Posterior localization of Aub and Vas was lost in oocytes of all *Mkrn1* mutant alleles (Figs 3M-T). Finally, Orb localization was unaffected in *Mkrn1*^*W*^ (Figs 3U and V), but was concentrated at the posterior in *Mkrn1*^*S*^ (Fig 3W). Many *Mkrn1*^*N*^ egg chambers included a single Orb-positive cell (Fig 3X), indicating that in these cases oocyte differentiation had taken place. Importantly, normal accumulation of all proteins could be restored by *nos*>GAL4 driven expression of a tagged *Mkrn1* transgene (Fig S5), confirming the specificity of the *Mkrn1* phenotypes.

**Figure 3.**
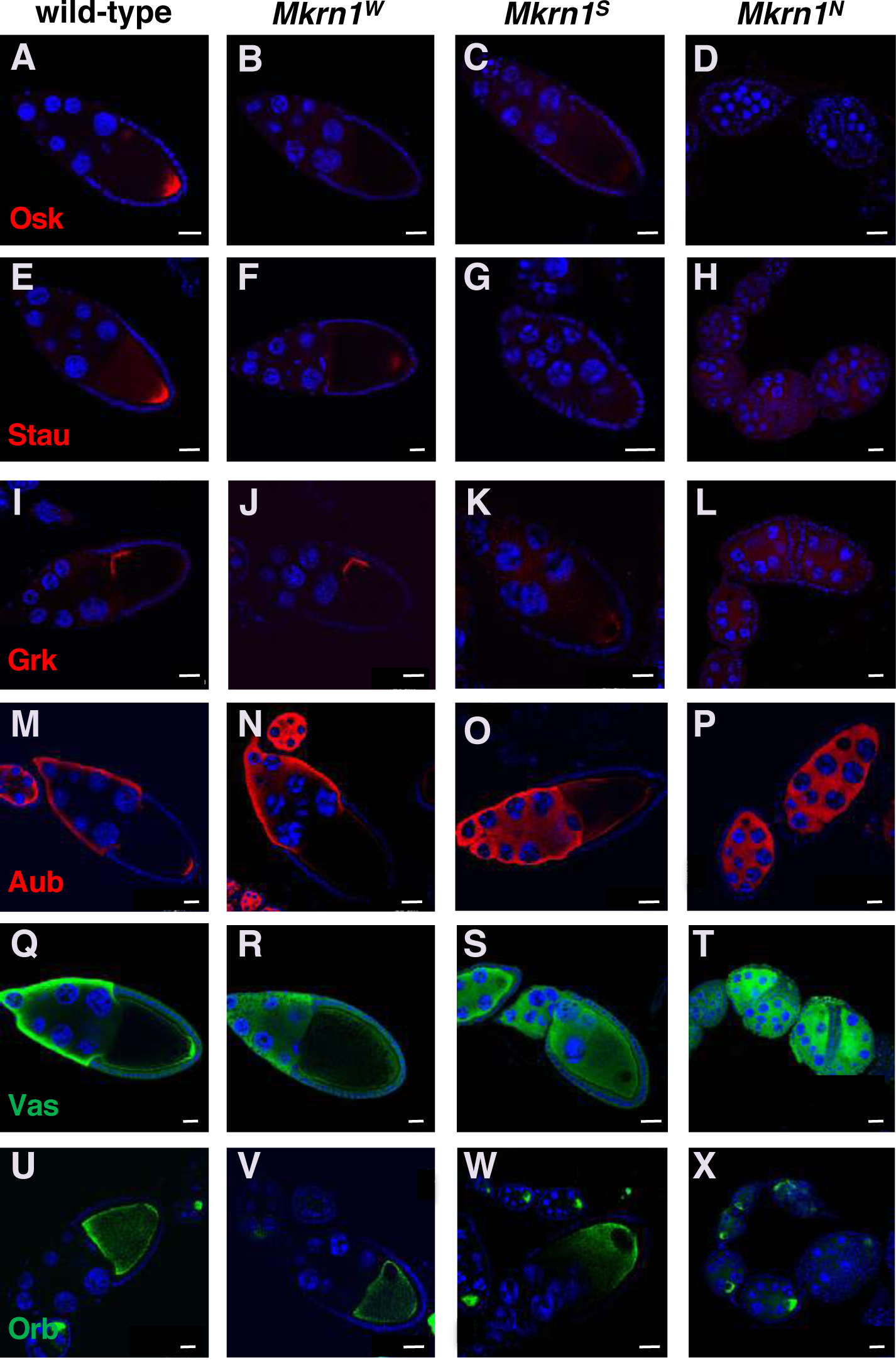
Mkrn1 mutations affect accumulation of proteins involved in axis patterning. (**A**-**D**) Posterior accumulation of Osk is greatly reduced in stage 10 *Mkrn1*^*W*^ and *Mkrn1*^*S*^ oocytes as compared with wild-type. Osk is nearly undetectable in *Mkrn1*^*N*^ egg chambers. Scale bars, 25 μm. (**E**-**H**) Posterior accumulation of Stau is greatly reduced in stage 10 *Mkrn1*^*W*^ and *Mkrn1*^*S*^ oocytes as compared with wild-type. Stau is nearly undetectable in *Mkrn1*^*N*^ egg chambers. Scale bars, 25 μm. (**I**-**L**) Anterodorsal accumulation of Grk is normal in stage 10 *Mkrn1*^*W*^ oocytes. Grk remains associated with the oocyte nucleus and is mislocalized to the posterior in stage 10 *Mkrn1*^*S*^ oocytes. Grk is present at uniformly low levels or undetectable levels in all germ cells in *Mkrn1*^*N*^ egg chambers. Scale bars, (**I**-**K**) 20 μm, (**L**) 25 μm. (**M**-**P**) Posterior accumulation of Aub is greatly reduced in stage 10 *Mkrn1*^*W*^ and *Mkrn1*^*S*^ oocytes as compared with wild-type. Aub is present at uniform levels in all germ cells in *Mkrn1*^*N*^ egg chambers. Scale bars, 20 μm. (**Q**-**T**) Posterior accumulation of Vas is greatly reduced in stage 10 *Mkrn1*^*W*^ and *Mkrn1*^*S*^ oocytes as compared with wild-type. Vas is present at uniform levels in all germ cells in *Mkrn1*^*N*^ egg chambers. Scale bars, 25 μm. (**U**-**X**) Accumulation of Orb is similar in wild-type and *Mkrn1*^*W*^ oocytes, but Orb is more concentrated in the posterior of *Mkrn1*^*S*^ oocytes. In early-stage *Mkrn1*^*N*^ egg chambers there is usually a single Orb-positive cell, indicating that some steps toward oocyte differentiation are able to take place. Scale bars, (**U**-**W**) 20 μm, (**X**) 25 μm.

### *Mkrn1*^*W*^ oocytes transiently accumulate *osk* mRNA at the posterior pole but do not produce Osk protein

To investigate whether the primary effect of *Mkrn1*^*W*^ on Osk deployment occurred at the level of RNA localization or translation, we examined more closely the effects of *Mkrn1*^*W*^ on *osk* mRNA localization and Osk translation using *in situ* hybridization for *osk* mRNA and immunostaining for Osk on the same samples (Fig 4A). In stage 7 egg chambers *osk* mRNA accumulation is robust in both wild-type and *Mkrn1*^*W*^ oocytes. Faint expression of Osk is occasionally visible in wild-type stage 7 oocytes but never in similarly staged *Mkrn1*^*W*^ oocytes. This difference becomes more pronounced in stages 9 and 10A. While 95% (41/43) of stage 9-10A *Mkrn1*^*W*^ oocytes show posterior accumulation of *osk* mRNA, only 7% (3/43) of *Mkrn1*^*W*^ oocytes show even a faint posterior signal for Osk protein. All (20/20) similarly-staged wild-type oocytes show strong posterior accumulation of both *osk* mRNA and Osk protein (Fig 4A). Subsequently, posterior accumulation of *osk* mRNA is lost in *Mkrn*^*W*^ oocytes. In 88% of stage 10B *Mkrn1*^*W*^ oocytes (30/34), *osk* mRNA remains in a tight focus, but is no longer anchored at the posterior pole (Fig 4A). *osk* translation remains repressed with only 6% of stage 10B *Mkrn1*^*W*^ oocytes (2/34) having detectable posterior Osk. In later *Mkrn1*^*W*^ oocytes neither posterior *osk* mRNA nor Osk protein are detectable. We conclude from these experiments that Mkrn1 is not directly required for the localization of *osk* mRNA at the posterior pole but is necessary for its translation. In agreement with previous observations [9,26,27,39], posterior accumulation of *osk* mRNA is not maintained in *Mkrn1*^*W*^ oocytes likely due to the failure of accumulating Osk protein. To confirm these results, we compared Osk protein levels by western blot analysis and *osk* mRNA levels by quantitative PCR (qPCR). Consistent with our immunostainings, we observed a pronounced reduction in Osk protein (Fig 4B), but not of *osk* mRNA (Fig 4C), in all *Mkrn1* alleles.

**Figure 4.**
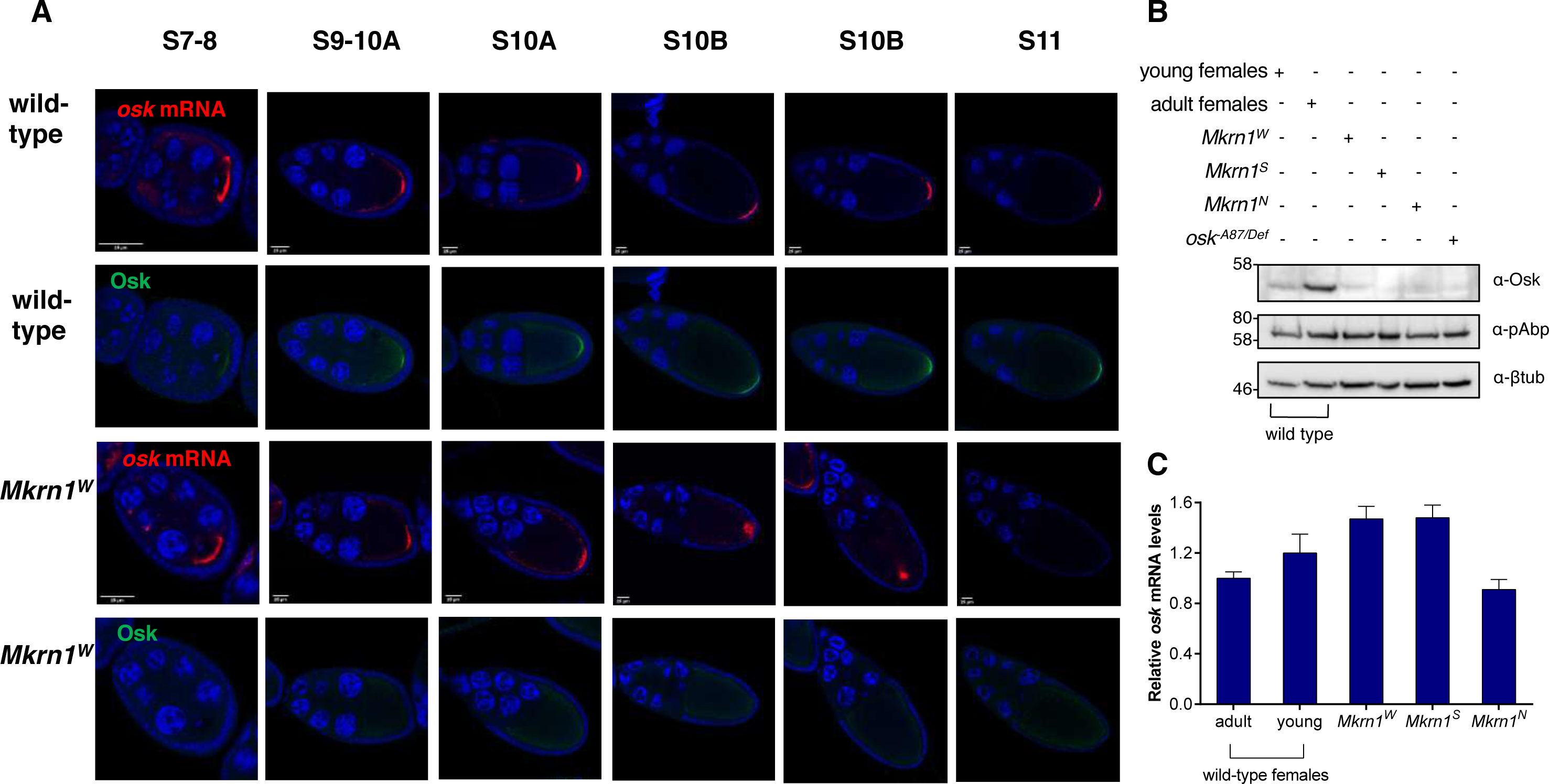
Translation of*osk* mRNAs is impaired in *Mkrn1*^*W*^ ovaries. (**A**) Fluorescent *in situ* hybridization for *osk* mRNA (red) with co-immunostaining for Osk protein (green) in wild-type and *Mkrn1*^*W*^ egg chambers. For each genotype and in each column the top and bottom images are of the same egg chamber. In wild-type oocytes posterior accumulation of *osk* mRNA and Osk protein is robust and stable from stage 9 onward. In *Mkrn1*^*W*^ oocytes accumulation of *osk* mRNA resembles the wild-type pattern through stage 10A but is not maintained, while Osk protein is rarely detectable at the oocyte posterior at any stage. Scale bars, 25 μm. (**B**) Western blot analysis from ovary lysates of various genotypes stained for Osk, pAbp and β-Tubulin. Osk protein levels are greatly reduced in all *Mkrn1* mutant alleles. 1-day-old young females have not yet completed oogenesis and were used as a control for *Mkrn1*^*S*^ and *Mkrn1*^*N*^ ovaries which also lack late-stage egg chambers, where Osk is most abundant. (**C**) RT-qPCR experiments measuring ovarian *osk* mRNA levels (normalized to *RpL15* mRNA) in the same genotypes as **(B)**. Error bars depict Stdev of two biological replicates.

### *Mkrn1* mutants affect localization of other maternal mRNAs

We used fluorescent *in situ* hybridization to investigate the distribution of several other mRNAs involved in patterning in *Mkrn1* mutants. Consistent with what we observed for Grk protein, localization of *grk* mRNA was similar to wild-type in *Mkrn1*^*W*^, but remained at the posterior in *Mkrn1*^*S*^ oocytes (Figs S6A-C). Posterior accumulation of *osk, nos* and *polar granule component* (*pgc*) mRNAs was also lost in *Mkrn1*^*W*^ embryos (Figs S6D-I).

### Mkrn1 physically associates with factors involved in *osk* mRNA regulation

To gain further insights into the molecular pathways underlying Mkrn1 function we sought to identify potential cofactors. For this purpose, we expressed Myc-tagged Mkrn1 in *Drosophila* S2R+ cultured cells and carried out immunoprecipitation (IP) experiments followed by mass spectrometry analysis. We also repeated this experiment using a version of Mkrn1 carrying a point mutation in the RING domain (Mkrn1^RING^), as we noticed that this construct was expressed at a much higher level compared to the wildtype one, which appears to be unstable after transfection in cells (Figs S7A-B). Similar stability characteristics have also been reported for mammalian MKRN1 [33]. Numerous RNA-binding proteins were enriched after IP, in particular when using Mkrn1^RING^ as bait (Figs S7C-D). Among those, several have been already linked to *osk* mRNA localization and translation [20,37,40-42]. To validate these interactions and address whether an RNA intermediate was involved, we performed co-IP experiments in S2R+ cells between Mkrn1^RING^ and various interaction partners in the presence or absence of RNase T1. Using this approach, we could confirm the interaction of Mkrn1^RING^ with polyA binding protein (pAbp), IGF-II mRNA-binding protein (Imp), eukaryotic initiation factor 4G (eIF4G), Squid (Sqd) and maternal expression at 31B (me31B), all in an RNA-independent manner (Fig S8). Interestingly, several of these components have already been shown to interact with each other [40,41,43-45]. We confirmed that interactions between FLAG-tagged Mkrn1 and pAbp as well as eIF4G also occur in ovaries (Fig 5A, Fig S8F).

**Figure 5.**
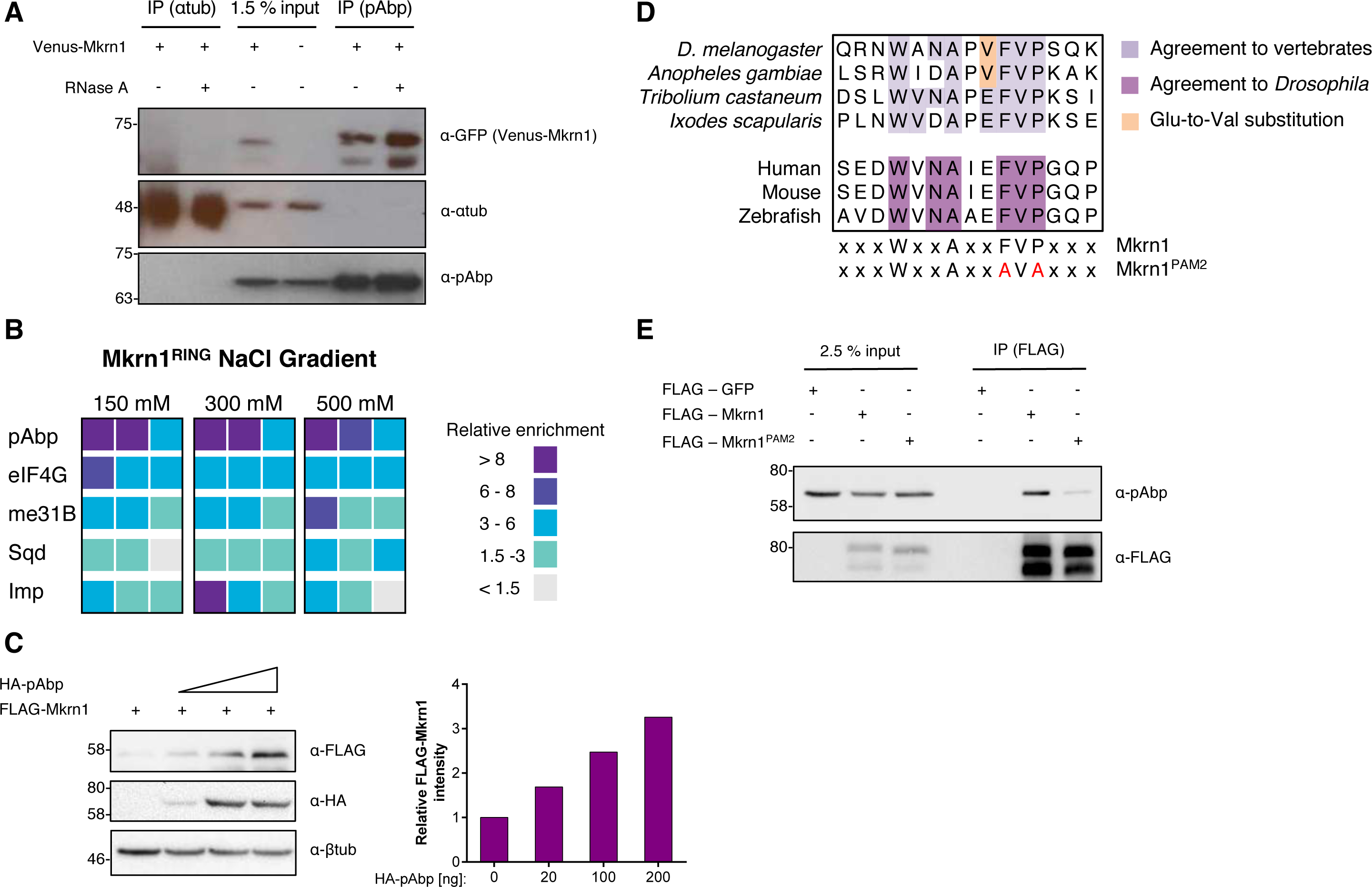
Mkrn1 interacts strongly with poly(A) binding protein. (**A**) Western blot analysis of co-IP experiments between Venus-Mkrn1 and pAbp. α-Tubulin (lanes 1, 2) and ovaries lacking the Venus-Mkrn1 transgene (lane 4) were used as negative controls. (**B**) Summary of co-IP experiments between Myc- or GFP-tagged interacting proteins and Mkrn1^RING^. The relative enrichment of Mkrn1^RING^ signal in each IP compared to controls are depicted for three individual experiments. (**C**) Co-expression of pAbp stabilizes Mkrn1. FLAG-Mkrn1 was co-transfected with increasing levels of HA-pAbp in S2R+ cells. Left: Proteins were examined using immunoblotting. Right: Intensities of FLAG-Mkrn1 levels were quantified and normalized to intensities of β-tubulin. The relative intensity was normalized to *Mkrn1* mRNA levels (normalized to *rpl15* mRNA) analyzed by qPCR. (**D**) PAM2 motif alignment in different species. Comparison between *Drosophila* and human PAM2 motif revealed a Glu to Val substitution (orange) in the consensus sequence. The conserved amino acid sequence to *Drosophila* (dark purple) is indicated below. The PAM2 motif was mutated using two amino acid substitutions at positions 90 and 92 to alanine (F90A and P92A). (**E**) Western blot analysis of co-IP experiments of FLAG-Mkrn1 and pAbp. The interaction of pAbp and Mkrn1 is reduced when the PAM2 motif is mutated.

To further address the strength of these interactions, co-IPs were repeated in a variety of salt concentrations. In this way we found that pAbp and eIF4G interact most strongly with Mkrn1 and these interactions were maintained upon stringent washes (Fig 5B, Fig S9). Consistent with this result, the stability of Mkrn1 itself was enhanced upon co-transfection of pAbp in S2R+ cells (Fig 5C). Mammalian MKRN1 contains a PCI/PINT associated module 2 (PAM2) motif, that is present in several pAbp binding proteins and serves in vertebrate MKRN1 for binding to PABP [36]. We identified a similar motif in *Drosophila* Mkrn1 (Fig 5D), but with one variation compared to human (V instead of E at position 9) that likely explains why this PAM2 motif was not recognized previously. To address the functionality of this motif we repeated the co-IPs, and found that when this domain was mutated (Mkrn1^PAM2^, Fig 5D), the interaction between Mkrn1 and pAbp was compromised (Fig 5E). Based on these data we conclude that Mkrn1 exists in one or several complexes that contain factors involved in the regulation of *osk* mRNA translation, and stably interacts with pAbp via its PAM2 motif.

### Mkrn1 associates specifically with *osk* mRNA *in vivo*

The effects we observed on *osk* translation in late *Mkrn1* egg chambers prompted us to test whether Mkrn1 can interact with *osk* mRNA. To assess whether Mkrn1 can bind to RNA in general, we immunoprecipitated FLAG-Mkrn1 after transfection of S2R+ cells and UV crosslinking. Mkrn1-bound RNA was subsequently labeled, and the protein-RNA complexes were visualized by autoradiography (Fig S10A). While a higher concentration of RNase I (1/50 dilution) resulted in a focused band, a lower concentration (1/5000 dilution) produced a shift of the Mkrn1-RNA complexes, demonstrating the RNA binding ability of *Drosophila* Mkrn1. We next repeated this experiment with various mutations in different Mkrn1 domains (Fig S10A). While mutations that alter the RING (Mkrn1^RING^) or the ZnF2 domain (Mkrn1^ZnF2^) behave as wild-type Mkrn1, deletion of the ZnF1 domain (Mkrn1^ΔZnF1^) resulted in a reduction of labeled Mkrn1-RNA complexes. These findings demonstrate that the ZnF1 domain is critical for association of Mkrn1 with RNA.

To address whether Mkrn1 associates with specific mRNAs *in vivo*, we overexpressed either wild-type Mkrn1 or Mkrn1^ΔZnF1^ in ovaries and performed RNA IP (RIP) experiments. The enrichment of different mRNAs was analyzed by qPCR using primers that bind to the 3’ UTRs of the respective transcripts. Interestingly, we observed that *osk* mRNA was substantially enriched in Mkrn1 IPs, but much less so when using Mkrn1^ΔZnF1^(Fig 6A, Fig S10B). On the other hand, *bcd* and *grk* mRNAs were not detected above background levels in either RIP experiment. This provides evidence that Mkrn1 binds specifically to *osk* mRNA *in vivo* and that its ZnF1 domain is important for this interaction.

**Figure 6.**
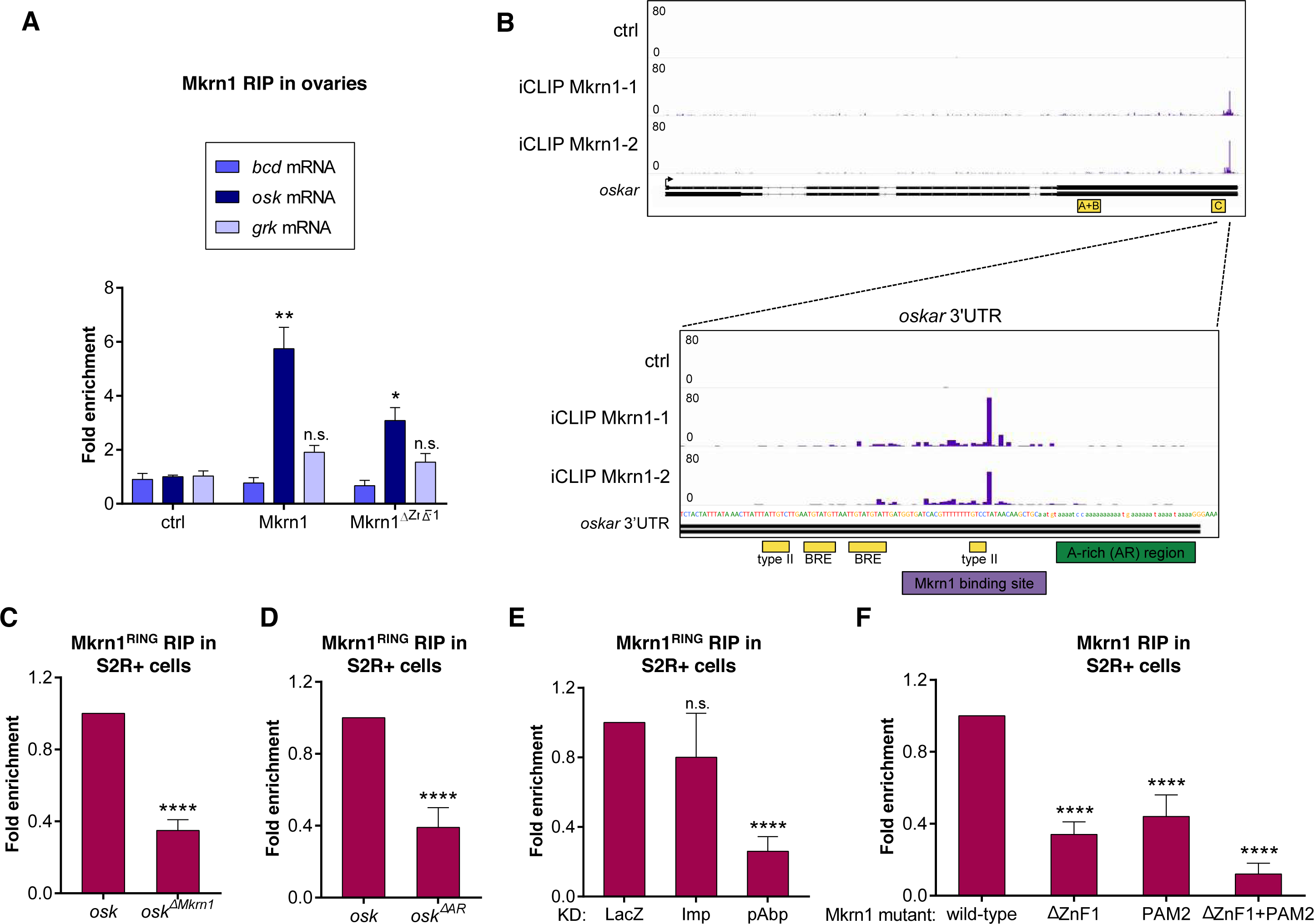
Mkrn1 associates specifically with the 3’ UTR of *osk* mRNA. **(A)** RIP experiment. Either FLAG-tagged Mkrn1 or Mkrn1^ΔZnF1^ was overexpressed in *Mkrn1*^*N*^ ovaries using *nos>GAL4* driver. Enrichment of different transcripts was analyzed by RT-qPCR using primers that bind to the respective 3’ UTRs. Error bars depict SEM, n=3. Multiple t-test was used to analyze significant changes compared to control IP. **(B)** iCLIP results showing specific binding of Mkrn1 to *osk* in a region of the 3’ UTR that partially overlaps with the BRE-C (yellow). The peaks indicate the crosslinking events of Mkrn1 to *osk* 3’ UTR (purple). The Mkrn1 binding site is upstream of an A-rich sequence (AR, green). Data of two technical replicates for FLAG-Mkrn1 are shown. **(C-E)** RIP experiments of FLAG-Mkrn1^RING^ in S2R+ cells. Enrichment of *luciferase*-*osk*-3’UTR transcript was analyzed by RT-qPCR. Relative change in binding compared to *luciferase*-*osk*-3’UTR or control knock down is depicted. Error bars depict SEM, n ≥ 3. (**C)** RIP experiment showing that Mkrn1^RING^ binding to *osk* 3’ UTR is compromised by a deletion of the Mkrn1 binding site (*osk*^Δ*Mkrn1*^, deletion of nucleotides 955-978 of *osk* 3’ UTR). **(D)** Binding of Mkrn1^RING^ to *osk* 3’ UTR is reduced when using a deletion of the A-rich region of *osk* 3’ UTR (*osk*^Δ*AR*^, deletion of nucleotides 987-1019 of *osk* 3’ UTR) **(E)** RIP experiments were performed in either control cells or upon depletion of Imp or pAbp. Depletion of pAbp compromises binding of Mkrn1. **(F)** RIP experiments showing that FLAG-Mkrn1 binding to *luciferase*-*osk*-3’ UTR is dependent on the ZnF1 domain as well as on the PAM2 motif. Enrichment was analyzed using RT-qPCR and relative change in binding compared to RIP of FLAG-Mkrn1 is illustrated. Error bars depict SEM, n ≥ 3.

To determine precisely where Mkrn1 binds *osk* mRNA, we performed crosslinking and IP (iCLIP) experiments after transfection of a tagged *Mkrn1* construct in S2R+ cells (Fig S10C). Since *osk* is poorly expressed in these cells we co-transfected a genomic construct of *osk* under the control of an *actin* promoter. We found specific binding sites in a handful of genes, including *osk* (Fig S10D), in which Mkrn1 binding sites were located in the distal part of the 3’ UTR (Fig 6B). These sites fall just upstream of an A-rich sequence that associates with pAbp [37]. Moreover, the binding site of Mkrn1 partially overlaps with the BRE-C site, which is bound by Bru1 and is required for both repression and activation of *osk* translation [24]. To validate the identified Mkrn1 binding sites, we performed RIP experiments in S2R+ cells using different 3’ UTRs fused to the firefly luciferase coding sequence. We found that Mkrn1 binds strongly to *osk* 3’ UTR but not to *grk* 3’ UTR (Fig S11A). In contrast, deletion of the Mkrn1-bound site identified with iCLIP (*osk*^ΔMkrn1^, deletion of nucleotides 955-978 of *osk* 3’ UTR) greatly reduced the interaction of Mkrn1 to *osk* 3’ UTR (Fig 6C, Fig S11B).

As the Mkrn1 binding site lies just upstream of the A-rich region (AR) we wondered whether the AR would also have an impact on Mkrn1 binding. To test this possibility, we deleted the AR (*osk*^ΔAR^, deletion of nucleotides 987-1019 of *osk* 3’ UTR) and examined Mkrn1 binding in S2R+ cells. We observed a decrease of Mkrn1 binding similar to that observed when deleting the Mkrn1 binding sites (Fig 6D, Fig S11C). Thus, we conclude that the AR enhances Mkrn1 binding to *osk* 3’ UTR. As Mkrn1 forms a stable complex with pAbp, our results further suggest that pAbp binding to the AR stabilizes Mkrn1 and therefore enhances its interaction with *osk*. Accordingly, reducing pAbp levels by RNAi, but not the level of Imp, another Mkrn1 interactor, dramatically decreased Mkrn1 association with *osk* mRNA in *Drosophila* cultured cells (Fig 6E, Figs S11D and E). Mutation of the PAM2 domain that enables the interaction with pAbp also resulted in reduced binding to *osk* 3’ UTR (Fig 6F, Figs S11F). Consistent with these results, mutating both the ZnF1 domain and the PAM2 motif led to almost complete loss of binding to *osk* 3’ UTR. Collectively, our results indicate that Mkrn1 binds specifically to the 3’ end of *osk* 3’ UTR via its ZnF1 domain and this association is further stabilized through the interaction with pAbp.

### Mkrn1 competes with Bru1 for binding to *osk* 3’ UTR

Our observation that Mkrn1 binds to the BRE-C prompted us to test whether Mkrn1 and Bru1 may compete for binding to *osk* 3’ UTR. To this end, we first examined whether we can recapitulate Bru1 binding to *osk* mRNA in S2R+ cells. As Bru1 is normally not expressed in this cell type, cells were co-transfected with a *Bru1* construct tagged with GFP along with the *luciferase*-*osk*-3’UTR reporter. RIP experiments confirmed previous findings that Bru1 strongly binds to *osk* 3’ UTR (Fig S12A, [18]). We next repeated this experiment upon knockdown of *Mkrn1* mRNA. Strikingly, while the overall level of Bru1 was not affected (Fig S12B), its binding to *osk* 3’ UTR was significantly increased (Fig 7A, Figs S12B and C). As pAbp is required to stabilize Mkrn1 binding to *osk* 3’ UTR, we wondered whether this interaction is necessary for modulating Bru1 binding. Indeed, knockdown of pAbp resulted in an increase in Bru1 binding to *osk* 3’ UTR as well (Fig 7B, Fig S12D). Next, to assess whether Mkrn1 competes with Bru1 for binding to the *osk* 3’ UTR *in vivo*, we repeated RIP experiments using ovarian extracts. Bru1 binding was assessed using an antibody directed against endogenous Bru1 and its association with *osk* mRNA was subsequently analyzed by qPCR. Similar to S2R+ cells, Bru1 binding to *osk* 3’ UTR was significantly increased in *Mkrn1*^*W*^ mutant ovaries (Fig 7C, Fig S12E). Thus, we conclude that Mkrn1 restricts Bru1 binding to *osk* 3’ UTR and this effect is enhanced by the interaction of Mkrn1 with pAbp.

**Figure 7.**
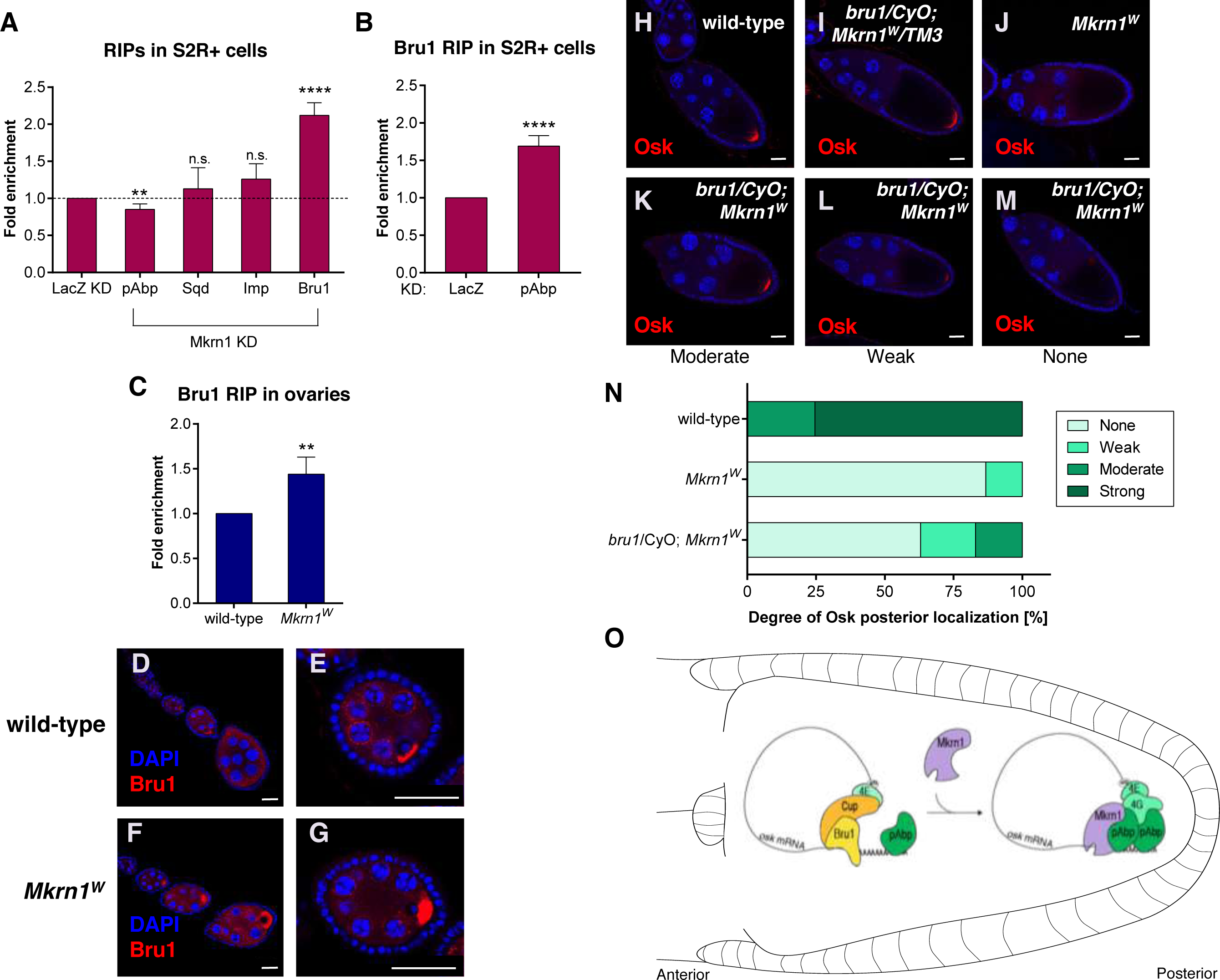
Mkrn1 competes with Bru1 for binding to *osk* mRNA. (**A**) RIP experiments in either control cells, or upon knockdown of Mkrn1. Binding of the indicated proteins to *luciferase*-*osk*-3’UTR was monitored by RT-qPCR. The relative fold change in recovered RNA upon Mkrn1 knockdown is illustrated. Error bars depict SEM, n=4. (**B**) Bru1 binding to *luciferase*-*osk*-3’UTR upon *pAbp* knockdown was analyzed using RT-qPCR. The relative fold change in binding of GFP-Bru1 to *luciferase*-*osk*-3’UTR compared to control knockdown is shown. Error bars depict SEM, n=3. (**C**) RIP experiments in either control (ctrl) or *Mkrn1*^*W*^ ovaries using α-Bru1 antibody. The relative fold change in recovered endogenous *osk* RNA compared to *Mkrn1*^*W*^ ovaries is shown. Error bars depict SEM, n=3. (**D**-**G**) Immunostaining experiments showing Bru1 distribution in (**D**-**E**) wild-type and (**F**-**G**) *Mkrn1*^*W*^ early-stage egg chambers. Note the more prominent accumulation of Bru1 in the oocyte in the *Mkrn1* mutant. Scale bars, (**D** and **F**) 25 μm; (**E** and **G**) 20 μm. (**H**-**M**) Stage-10 egg chambers of the genotypes indicated immunostained with α-Osk. Posterior accumulation of Osk is restored to a variable degree (**K**-**M**) in *Mkrn1*^*W*^ oocytes when heterozygous for *bru1*. Scale bars, 25 μm. (**N**) Quantification of posterior Osk localization in oocytes depicted in **(H-M)**. (**O**) Model depicting activation of *osk* translation via Mkrn1. Mkrn1 and pAbp are recruited to the *osk* 3’ UTR, displacing Bru1 and promoting translation initiation at the posterior pole.

To further address the relationship between Mkrn1 and Bru1, we examined whether the *Mkrn1*^*W*^ mutation affects Bru accumulation during oogenesis. In wild-type ovaries, Bru1 is expressed in all germline cells and accumulates to a modest degree in the oocyte during early oogenesis, (Fig 7D-E, [19]). In contrast, in *Mkrn1*^*W*^ ovaries, oocyte accumulation of Bru1 during early stages was much more pronounced (Fig 7F-G). As *osk* mRNA accumulates in early oocytes, this is consistent with Bru1 having an increased binding affinity for *osk* in the absence of Mkrn1. Nevertheless, this is not definitive as Bru1 can accumulate in the early oocyte in the absence of *osk* mRNA [46].

If Mkrn1 activates *osk* translation by displacing Bru1, we would predict that lowering *bru1* genetic dosage should suppress the *Mkrn1* phenotype. To test this hypothesis, we used a strong *bru1* allele (*bru1*^*QB*^, [19,47]). Indeed, removing one copy of *bru1* was sufficient to substantially rescue Osk protein level and posterior localization in *Mkrn1*^*W*^ female oocytes as measured by immunostainings (Figs 7H-N). We also observed a higher survival rate of embryos produced from *Mkrn1*^*W*^ females that were heterozygous for *bru1*^*QB*^ as compared to controls (620/2058, 30.1% vs 40/1222, 3.3%). Taken together, these experiments support a model in which Mkrn1 activates *osk* translation by displacing Bru1 binding to the *osk* 3’ UTR (Fig 7O).

## Discussion

Our data indicates that Mkrn1 is essential for embryonic patterning and germ cell specification. By taking advantage of a new allele that specifically disrupts Mkrn1 binding to RNA, we demonstrate that Mkrn1 exerts this function primarily via regulating *osk* translation by antagonizing Bru1 binding.

Control of *osk* translation has been studied in depth, revealing a complex spatio-temporal interplay between repressing and activating factors [48]. Relief of translational repression and activation of *osk* translation is likely to involve multiple redundant mechanisms. For example, Bru1 can be phosphorylated on several residues, and phosphomimetic mutations in these residues inhibit Cup binding in pulldown assays. However, these do not seem to affect translational repression activity *in vivo* [23]. In agreement with this, we did not observe a change in Bru1 binding to itself nor to Cup upon depletion of Mkrn1 (data not shown). Stau, Aub, Orb and pAbp have also been implicated in activating *osk* translation [14,28,37,49]. It is unlikely that Mkrn1 controls *osk* translation by recruiting Stau as Stau still colocalizes with *osk* mRNA in *Mkrn1*^*W*^ oocytes (Figs S12F-H). Instead, we propose that Mkrn1 exerts its positive activity by competing with Bru1 binding on *osk* 3’UTR. This is evidenced by the overlap of their binding sites, the increased Bru1 binding to *osk* RNA upon Mkrn1 knockdown and by our observation that reducing *bru1* dosage is sufficient to partially alleviate *osk* translational repression.

Two distinct Bru1 binding regions (AB and C) are present in the *osk* 3’ UTR and are required for translational repression. However, the C region has an additional function in translational activation. Indeed, it was hypothesized that an activator binds the C region to relieve translational repression [24]. This activator was proposed to either be Bru1 itself, or a different protein that can bind both the BRE and the type II Bru1-binding site, which is what we observe for Mkrn1 (Fig 6B). Later work showed that Bicoid Stability Factor (BSF) binds the C region *in vitro* at the 3’ type II Bru1-binding site [50], which is also where we mapped Mkrn1 binding. Deletion of this site impacts embryonic patterning, yet depletion of BSF has no effect on Osk protein expression up to stage 10, indicating that initial activation of *osk* translation is effective even in the absence of BSF [50]. In this case, only late stage oocytes display reduced Osk accumulation. Therefore, it is possible that a concerted action of Mkrn1 and BSF exists at the type II site to trigger *osk* translation and sustain it at later stages.

The binding of Mkrn1 to mRNA seems to be extremely specific. We found that the binding to *osk* is dependent on a downstream A-rich sequence and on interaction with pAbp. Relevant to this, Bru1 binds to *grk* 3’ UTR in addition to *osk* [51,52], and several proteins that associate with Mkrn1 also associate with *grk* mRNA [43,44]. However, we found that Mkrn1 does not bind strongly to *grk*, which lacks poly(A) stretches in the proximity of its Bru1 binding sites, and consistently, we did not observe a regulatory role of Mkrn1 on Grk translation.

In addition to pAbp, it is noteworthy that Mkrn1 associates with other proteins previously implicated in *osk* localization and translational activation. Its interaction with eIF4G would be consistent with a role in alleviating Cup-mediated repression, as it could recruit eIF4G to the cap-binding complex at the expense of Cup. However, we did not observe an interaction between Mkrn1 and eIF4E (data not shown). The association between Mkrn1 and Imp is also intriguing as the *osk* 3’ UTR contains 13 copies of a five-nucleotide motif that binds to Imp [27]. This region is essential for *osk* translation but Osk accumulation is unaffected in *Imp* mutants, suggesting the involvement of another factor that binds these motifs [25,27,43]. In contrast to pAbp, we did not observe alteration of Mkrn1 binding when Imp was depleted, indicating that Imp is not required to stabilize Mkrn1 on *osk* mRNA.

The molecular links we uncovered between Mkrn1 and RNA-dependent processes in *Drosophila* are consistent with recent high-throughput analysis of mammalian MKRN1 interacting proteins [35]. RNA binding proteins, including PABPC1, PABPC4, and eIF4G1, were highly enriched among the interactors. In the same study, MKRN1 was also shown to interact with RNA. In addition, the short isoform of rat MKRN1 was shown to activate translation but the underlying mechanism remained unknown [36]. Since in vertebrates *MKRN* genes are also highly expressed in gonads and early embryos, it is possible that similar molecular mechanisms are employed to regulate gene expression at these stages [32]. Consistent with this, MKRN2 was recently found to be essential for male fertility in mice [53]. Thus, our study provides a mechanism that explains the role of Mkrn1 in translation and constitutes a solid framework for future investigations deciphering the role of vertebrates MKRN in post-transcriptional control of gene expression during gametogenesis and early development.

## Materials and Methods

### Generation of *Mkrn1* mutants using CRISPR/Cas9

The guide RNAs used were cloned into expression vector pDFD3-dU63gRNA (Addgene) according to manufacturer’s instructions. Different guide RNAs were used either alone (gRNA1 starting at nucleotide 64 of *Mkrn1* CDS and gRNA2 starting at nucleotide 363 of *Mkrn1* CDS) or in combination (gRNA3 starting at 387 nt of *Mkrn1* gene and gRNA4 starting at position 2239 nt). *vas*-Cas9 *Drosophila* embryos were injected with the purified plasmids containing the gRNA (500 ng/µl in H_2_O) and allowed to develop to adulthood. Each male was crossed with double balancer females. Genomic PCR from single flies was prepared and tested for CRISPR/Cas9 induced mutations using the T7 endo I (BioLabs) assay or by PCR using primers that bind in proximity to the guide RNA targeting site. A list of gRNAs as well as primers is appended (Table S3).

**Table S3.**
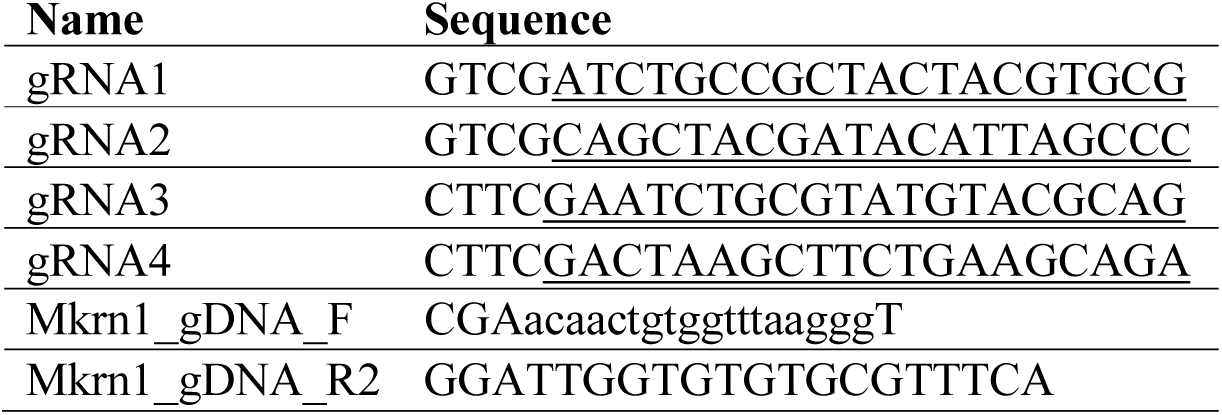
List of gRNAs used in this study and primers used to validate mutations of the *Mkrn1* gene by CRISPR/ Cas9.

### Immunostaining and confocal imaging

Ovaries were dissected and fixed in 4% paraformaldehyde in PBS for 20 min at room temperature (RT). After 4-5 washes in PBST (PBS containing 0.3% Triton-X100) ovaries were permeabilized with 1% freshly prepared Triton-X100 in PBS for 1 h. The ovaries were blocked in 2% BSA/PBST overnight. Dispersed egg chambers were then incubated with the primary antibodies diluted in 2% BSA/PBST at RT for 4 h or overnight at 4°C. The washed egg chambers were incubated with conjugated secondary antibodies at 1:500 at RT for 4 h or overnight at 4°C. DAPI (1 ng/ml) was added in the last wash to counter-stain the nuclei for 30 min. After 2-3 washes with PBST the mounting medium containing 1% DABCO was added and the samples were equilibrated for 30 min or overnight. The stained samples were mounted on glass slides and sealed with nail varnish for microscopy imaging. Rabbit polyclonal antibodies against Vas, Osk, Aub, Grk were generated in the Lasko lab; rabbit α-Stau is from the St Johnston lab; rabbit α-Bru1 is from the Ephrussi lab and rabbit α-pAbp is from the Sonenberg lab; mouse monoclonal antibodies against Orb, Sqd, and Lamin were purchased from the Developmental Studies Hybridoma Bank; mouse α-GFP and rabbit α-Flag were purchased from Abcam and Sigma. Alexa Fluor 488 or 555 conjugated secondary antibodies were purchased from Molecular Probes, and pre-absorbed with fixed and blocked wild type ovaries to reduce background. Stained egg chambers were examined using a confocal microscope (Leica). Images were taken under 40 x oil lens by laser scanning and processed with ImageJ.

### *In situ* hybridization of embryos and ovaries and RNA-protein double labeling

cDNAs were used as templates for PCR to generate an amplified gene fragment with promoter sequences on each end. PCR products were purified via agarose gel extraction and used for *in vitro* transcription to generate digoxigenin-labeled RNA antisense probes with MAXIscript kit (Ambion). The length of each probe was about 1000 nt. *In situ* hybridization experiments were performed as described [54], using biotinylated α-DIG antibody and streptavidin-HRP followed by tyramide conjugation for development of FISH signal. For RNA-protein double labeling, ovaries or embryos were incubated in primary antibody against the protein of interest along with biotinylated α-DIG antibody at 4°C overnight. The tissue was washed, then detection reagent (fluorochrome-conjugated secondary antibody) along with streptavidin-HRP was added and incubated at 4°C overnight. Images were taken with confocal microscope (Leica).

### Embryo cuticle preparation and staining

Flies were transferred into egg-laying cages with apple juice agar plates and incubated at 25°C in the dark. Embryos were collected when 50-100 eggs had been laid and allowed to age for 24 h at 25°C. Embryos were collected in a sieve, dechorionated with 50% bleach for 2.5 min, washed with water, then transferred into PBST buffer (PBS + 0.1% Tween 20). For cuticle preparations, PBST buffer was removed, then 40-50 µl of Hoyer’s solution was added and embryos were kept at 4 °C overnight. Embryos in Hoyer’s solution were mounted on a glass slide, covered with a cover slip and incubated at 60-65 °C overnight. Dark-field images were taken with Leica DM6000B microscope.

### Staging

Staging experiment was performed as described [55] using *D*. *melanogaster w*^*1118*^ flies.

### Generation of transgenic flies

The constructs were made using Gateway technology (Invitrogen). Full-length wild-type *Mkrn1* cDNA was cloned into the pENTR entry vector using the pENTR/D-TOPO cloning kit (ThermoFisher). After verifying the sequence by PCR, the insert was subcloned into the expression vectors containing UASp promoter with different tags (pPVW and pPFW with Venus and FLAG tags at N-terminal respectively), and (pPWV and pPWF with Venus and FLAG tags at C-terminal respectively) by LR in vitro recombination. For Mkrn1^ΔZnF1^ pPFMW vector was used. Constructs were verified by sequencing and then injected into *yw* embryos. Progeny harboring the transgenes were crossed with double balancer flies to establish a variety of lines, and the insertion sites were mapped to either the second or third chromosome. Mkrn1 expression was then driven by crossing the transgenic lines with a *nos>Gal4* line (MTD). Expression of tagged Mkrn1 was verified by western blot analysis and immunostaining using anti-GFP (Abcam) or anti-FLAG (Sigma).

### Rescue experiment

Flies carrying Venus- or FLAG-Mkrn1 on the second chromosome were crossed with the *nos>Gal4* driver lines in three different Mkrn1 mutant backgrounds (*Mkrn1*^*W*^, *Mkrn1*^*S*^, and *Mkrn1*^*N*^). Progeny were collected and separated into two groups: (1) *Venus-* or *FLAG-Mkrn1/ nos>Gal4; Mkrn1*^*W*^ ^*(S*^ or ^*N)*^*/ Mkrn1*^*W*^ ^*(S*^ or ^*N)*^ and (2) *nos>Gal4/CyO*; *Mkrn1*^*W*^ ^*(S*^ or ^*N)*^*/ Mkrn1*^*W*^ ^*(S*^ or ^*N)*^. To perform hatching tests the same number of flies from each group was fed with yeast butter on apple juice plates for 1 d, embryos were collected and incubated at 25°C for 48 h to allow completion of hatching. Hatched and unhatched embryos were counted for each group. The data from several tests in the same group were pooled and the hatching percentage was calculated.

### Cell line

Drosophila S2R+ are embryonic derived cells obtained from Drosophila Genomics Resource Center (DGRC, Flybase ID: FBtc0000150).

### Cell culture, RNAi, transfection

Drosophila S2R+ cells were grown in Schneider’s medium (Gibco) supplemented with 10% FBS (Sigma) and 1% Penicillin-Streptomycin (Sigma). For RNAi experiments, PCR templates for the dsRNA were prepared using T7 Megascript Kit (NEB). S2R+ cells were seeded at the density of 10^6^ cells/ml in serum-free medium and 15 µg/ml of dsRNA was added to the cells. After 6 h of cell starvation, serum supplemented medium was added to the cells. dsRNA treatment was repeated after 48 h and cells were collected 24 h after the last treatment. A list of primers used to create dsRNA templates by PCR is appended (Table S4). Effectene (Qiagen) was used to transfect vector constructs in all overexpression experiments following the manufacturer’s protocol.

**Table S4.**
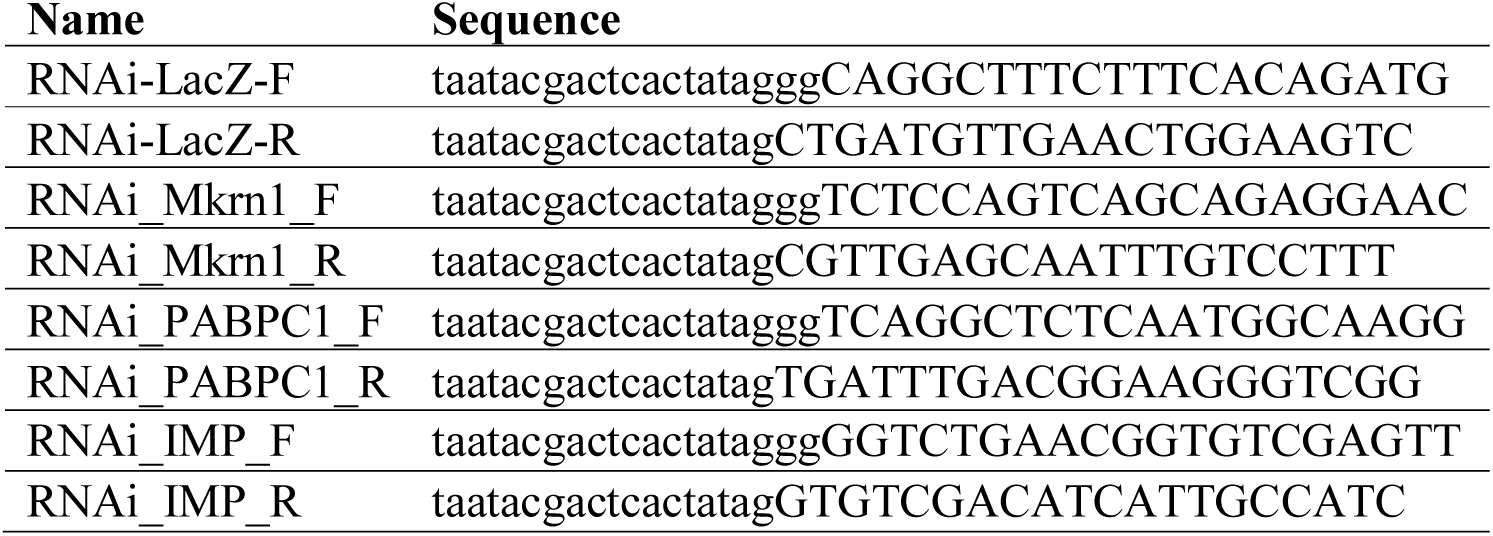
List of primers used to create PCR templates of dsRNAs.

### Immunoprecipitations (IPs)

For IP experiments in S2R+ cultured cells, protocol was followed as described [55] with minor changes: 2 mg of the protein lysates was incubated for 2 h with 10 µl of either Myc-Trap^®^ or GFP-Trap^®^ beads (Chromotek). To determine the dependence of interactions on RNA, 50 U of RNaseT1 (ThermoFisher) were added to the respective IP. To ensure the activity of RNase T1, lysates were incubated 10 min at RT prior to the incubation of lysate with antibody.

For IP experiments in ovaries, 150 µl of wet ovaries from 3-5 day old flies expressing Venus-Mkrn1 was homogenized on ice in 2 ml of cold IP buffer (1 X PBS, 0.4% Triton X-100, 1 mM MgCl_2_, 5% glycerol), containing protease inhibitors and PMSF. The extracts were diluted to 1.5 mg protein/ml. Each extract (0.66 ml) was mixed with 24 µg of anti-pAbp Fab antibody (Smibert lab, [56]), 17 µg of α-eIF4G rabbit antibody, or 15 µl of rabbit anti-α–Tubulin antibody (Abcam). When present, 100 µg RNase A (Qiagen) was added to the samples. Samples were incubated with rotation at 4 °C overnight, then mixed with 30 µl of protein A agarose beads (wet volume, Invitrogen) and incubated with rotation at RT for 1.5 h. The beads were washed three times with IP buffer. Bound material on the beads was eluted by boiling for 2 min in 40 µl of SDS loading buffer. 20 µl of the eluted sample, together with input samples, was used for western blot.

### RNA-Immunoprecipitation (RIP)

For RIP, S2R+ cells or ovaries were harvested and lysed in RIP buffer (150 mM NaCl, 50 mM Tris-HCl pH 7.5, 0.5% NP-40, 1 mM EDTA) supplemented with proteinase inhibitors (1.5 μg/ml Leupeptin, 1.5 μg/ml Pepstatin, 1.5 μg/ml Aprotinin and 1.5 mM PMSF) and RNase inhibitors (20 U/μl). S2R+ cells were lysed for 20 min at 4°C, subtracted to 2 cycles of sonication on a bioruptor (Diagenode) with 30 sec “ON”/“OFF” at low setting and the remaining cell debris was removed by centrifugation at 21,000 g for 10 min at 4°C. To remove lipids and cell debris, ovary lysates were centrifuged 4 times. Protein concentrations were determined using Bradford reagent (BioRad). 2 mg of protein lysate were incubated for 3 h with 2 μg of α-FLAG M2^®^ antibody (Sigma-Aldrich) pre-coupled to 20 μl of rotein G Dynabeads^®^ (Thermo Fisher Scientific) head-over-tail at 4°C. For RIP experiments analysing binding of Bru1 in ovaries, either 1 μl of rabbit α-Bru1 (gift from A. Ephrussi) or 2 μg of rabbit IgG (Millipore) were incubated with ovarian lysate over night at 4°C. 20 μl of protein G Dynabeads^®^ were added for 2 h after the incubation.

For every RIP experiment, beads were washed 4 x for 10 min in RIP buffer at 4°C. For immunoprecipitation of GFP-tagged Imp and Bru1 15 μl of GFP-Trap^®^ (Chromotek) were used. Lysates were prepared similar as above using RIPA buffer (140 mM NaCl, 50 mM Tris pH 7.5, 1 mM EDTA pH 8, 1% Triton X-100, 0.1% SDS, 0.1% sodium deoxycholate) supplemented with proteinase and RNase inhibitors. IP was performed for 2 h at 4°C and subsequently washed 4 x for 10 min with RIPA buffer.

RNA was eluted in TRIzol Reagent (ThermoFisher), 10 min at RT and subjected to RNA isolation and RT-qPCR. To obtain the depicted fold enrichment, individual transcripts were normalized to either *18S* or *RpL15*. Al least three biological replicates were performed for each experiment. If not stated differently statistical analysis was performed using one sample t-test.

To analyze IPs, 30% of beads were eluted in 1x SDS buffer (50 mM Tris pH 6.8, 2% SDS, 10% glycerol, 100 mM DTT, 0.05% Bromphenol Blue) at 95°C for 10 min. Eluted IP proteins were removed from the beads and analyzed by western blot together with input samples.

### Western blotting

Western blotting was performed as described [55]. Primary antibodies used were: mouse α-Myc 9E10 antibody (1:2000, Enzo Life Sciences); mouse α-FLAG M2^®^ antibody (1:1000, Sigma-Aldrich); rabbit α-GFP TP401 antibody (1:5000, Acris Antibodies); mouse α-HA F7 (1:1000, Santa-Cruz) rat α-HA (1:750, Roche); mouse α-β-Tubulin (1:5000, Covance), mouse α-α-Tubulin (1:20,000; Sigma), mouse α-GFP (1:500; Molecular probe), mouse α-ubiquitin (1:1000; Santa Cruz) Fab α-pAbp (2.5 µg in 5 ml), α-eIF4G rabbit antibody (1 µg in 5 ml), rabbit α-Osk (1:1000) antibody was a gift from A. Ephrussi.

### RNA isolation and measurement of RNA levels

Cells or tissues were harvested in TRIzol Reagent (ThermoFisher) and RNA was isolated according to the manufacturer’s instructions. DNA was removed with DNaseI treatment (NEB) and cDNA was prepared with M-MLV Reverse Transcriptase (Promega). The transcript levels were quantified using Power SYBR^®^ Green PCR Master Mix (ThermoFisher) using the indicated primer (Table S5).

**Table S5.**
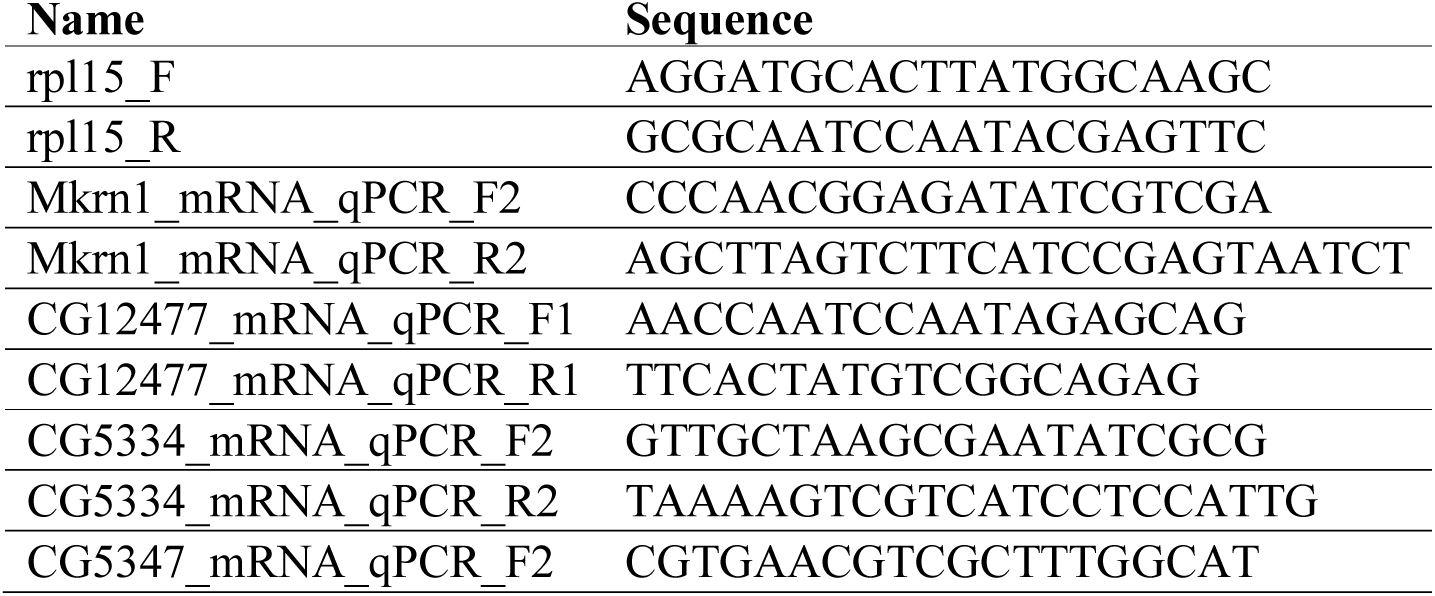
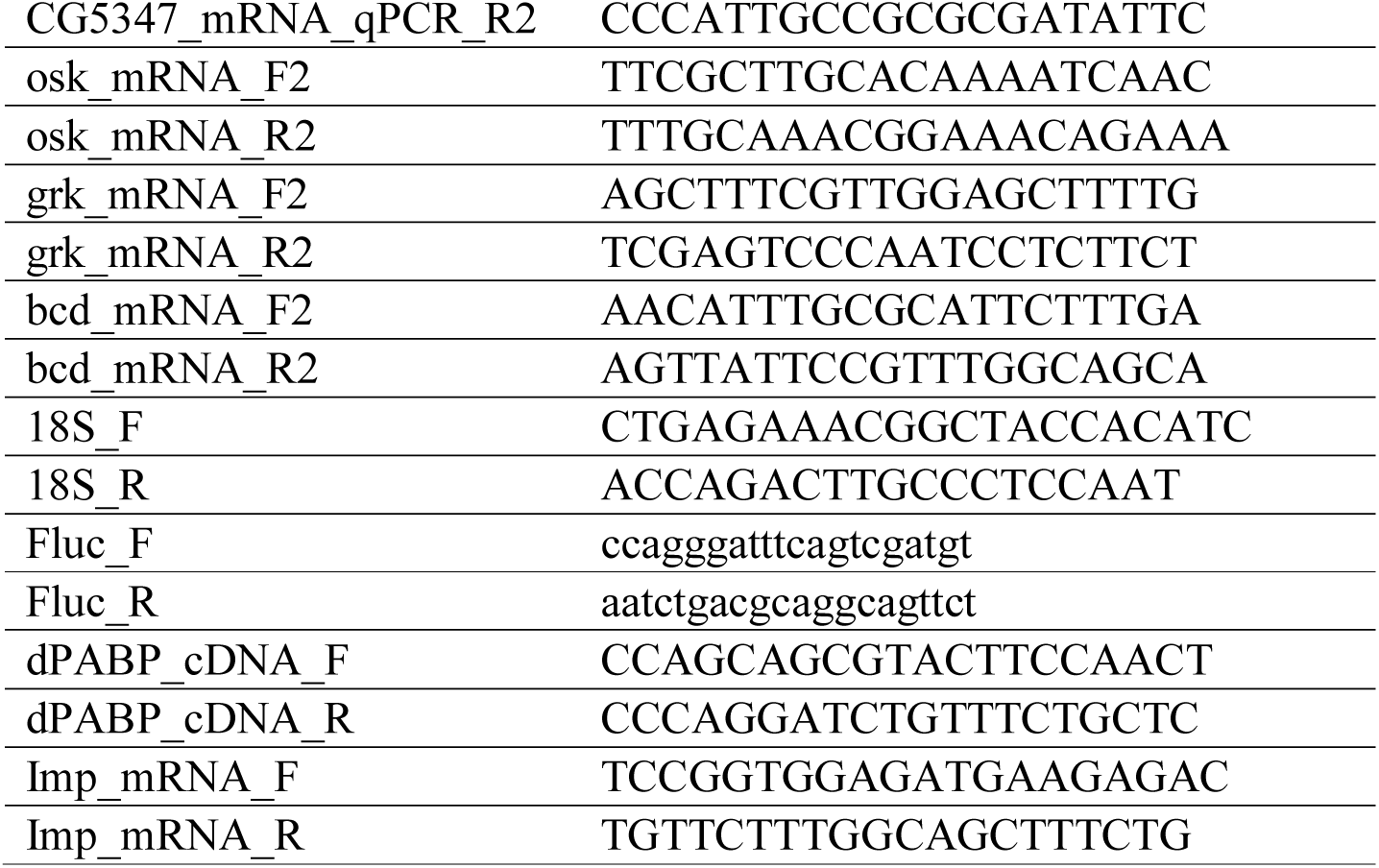
List of primers used to in qPCR experiments.

### LC-MS/MS

To identify binding partners of Mkrn1, either Myc-GFP as control or Myc-Mkrn1 were ectopically expressed in S2R+ cells. Upon lysis, Myc-GFP or Myc-Mkrn1 were immunoprecipitated as described above (see IP methods) with small adjustments: The IP buffer was additionally supplemented with 10 mM N-ethylmaleimide, 1 mM sodium orthovanadate, 5 mM β-glycerophosphate and 5 mM sodium fluoride. After IP, samples were eluted in 2x LDS buffer (Life Technologies) supplemented with 1 mM dithiothreitol for 10 min at 70°C and incubated with 5.5 mM 2-chloracetamide for 30 min at room temperature in the dark. All samples were prepared in parallel.

Conventional interactome analysis of the IP samples was performed as described before [57] with the following changes: The enriched proteins were separated by SDS-PAGE with a 4-12% Bis-Tris protein gel (NuPAGE, Thermo Scientific) and stained with Colloidal Blue Staining Kit (Life Technologies). Subsequently, proteins were in-gel digested using trypsin and digested peptides were then extracted from the gel. Concentration, clearance and acidification of peptides, mass spectrometry analysis, and peptide identification were performed as described before [57]. For peptide identification in MaxQuant (version 1.5.28), the DROME database from UniProtKB (release May 2016) was used. For label-free quantification (LFQ) at least 2 LFQ ratio counts (without fast LFQ) were activated.

The data table of LFQ values resulting from MaxQuant was filtered for potential contaminants, reverse binders and protein groups only identified by site. Furthermore, protein groups with less than two peptides and less than one unique peptide were also removed from further analysis. After log-transforming all remaining LFQ values, missing values were imputed by beta distributed random numbers between 0.1% and 1.5% of the lowest measured values. As a final filtering step, only protein groups having measured values for at least two replicates of at least one experimental condition were kept for further analysis. All filter and imputing steps were done with an in-house R script.

Differential protein abundance analysis was performed on log-transformed LFQ values between two conditions at the time using the R package limma (version 3.34.9, [58]). For each such comparison, only protein groups found in at least two replicates of at least one condition were kept and used. To visualize the interactome, the R package ggplot2 [59] was used. All protein groups with an FDR ≤ 0.05 and a log_2_ fold change of ≥ 2 were considered significantly changed.

The mass spectrometry proteomics data have been deposited to the ProteomeXchange Consortium via the PRIDE partner repository with the dataset identifier PXD011802.

### PAM2 motif alignment

Ortholog searches were performed using HaMStR-OneSeq [60]. Human MKRN1 and MKRN2 (UniProt identifiers: Q9UHC7, Q9H000) served as seed proteins and orthologs were searched within data from the Quest for Orthologs Consortium (release 2017_04, [61]). In order to identify functionally equivalent proteins, we calculated a unidirectional Feature Architecture Similarity (FAS) score that compares the domain architecture of the seed protein and the predicted ortholog [62]. Predicted orthologs with FAS < 0.7 were removed. The multiple sequence alignment of PAM2 motifs of makorin orthologs from selected arthropod and vertebrate species was generated using MAFFT v7.294b L-INS-i [63]. Since the PAM2 motif in all makorin proteins differs from the described consensus, a PAM2 hidden Markov model was trained on makorin PAM2 motifs and used for a HMMER scan (http://hmmer.org/) of the orthologs. Orthologs include species name, UniProt identifiers and amino acid (aa) positions of the PAM2 motif within the protein: *Drosophila melanogaster*, Q9VP20, 81-95 aa; *Anopheles gambiae*, Q7QF83, 57-71 aa; *Tribolium castaneum*, A0A139WP96, 159-173 aa; *Ixodes scapularis*, B7QIJ9, 119-133 aa; human, Q9UHC7, 163-177 aa; mouse, Q9QXP6, 163-177 aa; zebrafish, Q4VBT5, 120-134 aa.

### Individual-nucleotide resolution UV CrossLinking and ImmunoPrecipitation (iCLIP) and autoradiography

The iCLIP protocol was from [64] with the following adaptations: S2R+ cells were crosslinked with 150 mJ/cm^2^ of UV light and subsequently harvested. Cells were lysed in urea cracking buffer (50 mM Tris pH 7.5, 6 M urea, 1% SDS, 25% PBS) and sonicated using 2 cycles with 30 sec “ON”/“OFF” at low setting. Remaining cell debris was removed by centrifugation at 21,000 x g for 10 min at 4°C. Lysate was diluted 1:5 in IP buffer (150 mM NaCl, 50 mM Tris pH 7.5, 0.5% Tween-20, 0.1 mM EDTA) and incubated with 2 µg of anti-FLAG M2^®^ antibody (Sigma-Aldrich) pre-coupled to 40 µl of protein G Dynabeads^®^ (ThermoFisher Scientific) for 2 h at 4°C. The beads were washed 3x with high salt buffer and 3x with PNK buffer before Turbo DNase (Ambion) and RNase I (Ambion) treatment was performed. Beads were washed again 3x with high-salt buffer and 3x with PNK buffer to remove residual enzymes.

To analyze binding of Mkrn1 to RNA, RNA-protein complexes were analyzed using autoradiography using the same protocol as described above.

For high-throughput sequencing, libraries of 6 technical replicates for FLAG-Mkrn1 and 1 replicate for FLAG-GFP (ctrl) were prepared as described [65]. Multiplexed iCLIP libraries were sequenced as 75-nt single-end reads on an Illumina MiSeq sequencing system.

Sequencing qualities were checked for all reads using FastQC (version 0.11.5) (https://www.bioinformatics.babraham.ac.uk/projects/fastqc/). Afterwards, reads were filtered based on sequencing qualities (Phred score) of the barcode region. Reads with more than one position with a Phred score < 20 in the experimental barcode (positions 4 to 7 of the reads) or any position with a Phred score < 17 in the random barcode (positions 1 to 3 and 8 to 9) were excluded from subsequent analysis. Remaining reads were de-multiplexed based on the experimental barcode (positions 4 to 7) using Flexbar (version 3.0.0, [66]) without allowing any mismatch.

All following steps of the analysis were performed on the individual samples after de-multiplexing. Remaining adapter sequences were removed from the read ends using Flexbar (version 3.0.0) with a maximal error rate of 0.1 and a minimal overlap of 1 nt between the beginning of the adapter and the end of the read. Following adapter trimming, the first 9 nt of each read containing the experimental and random barcodes were trimmed off and added to the read name in the fastq files in order to keep this information for downstream analysis. Reads shorter than 15 nt were removed from further analysis.

Trimmed and filtered reads were mapped to the *Drosophila melanogaster* genome (Ensembl genome assembly version BDGP6) and its annotation (Ensembl release 90, [67]) using STAR (version 2.5.2b, [68]). When running STAR, up to two mismatches were allowed, soft-clipping was prohibited and only uniquely mapping reads were kept for further analysis. For further analysis, only unspliced reads were kept and analyzed.

Following mapping, duplicate reads were marked using the dedup function of bamUtil (version 1.0.13), which defines duplicates as reads whose 5’ ends map to the same position in the genome (https://github.com/statgen/bamUtil). Subsequently, marked duplicates with identical random barcodes were removed since they are considered technical duplicates, while biological duplicates showing unequal random barcodes were kept.

Resulting bam files were sorted and indexed using SAMtools (version 1.3.1, [69]). Afterwards, bedgraph files were created based on bam files, using bamToBed of the BEDTools suite (version 2.25.0; [70]), considering only the position upstream of the 5’ mapping position of the read, since this nucleotide is considered as the crosslinked nucleotide. Using bedGraphToBigWig of the UCSC tool suite [71], all bedgraph files were converted into bigWig files. The counts of all Mkrn1 iCLIP experiments were combined in one file.

In order to estimate binding site strength and to facilitate comparisons between binding sites (Fig S10D) we corrected for transcript abundance by representing the crosslink events within a binding site as a ‘signal-over-background’ ratio (SOB). The respective background was calculated as the sum of crosslink events outside of binding sites (plus 5 nt to either side) by the merged length of all exons. 3’ UTR lengths were restricted to 10 nt past the last Mkrn1 binding site or 500 nt if no binding site was present. SOB calculations were performed separately for each replicate and then averaged. No SOB value was assigned for ribosomal genes and genes with a background of < 10 crosslink events, resulting in SOB values for 184 binding sites in 46 targets.

The iCLIP data has been deposited to the NCBI’s Gene Expression Omnibus [72] and are accessible through GEO Series accession number GSE123052 (https://www.ncbi.nlm.nih.gov/geo/query/acc.cgi?acc=GSE123052).

## Cloning

To overexpress tagged proteins, the respective coding sequences were amplified and cloned into Gateway plasmids using *Asc*I and *Not*I restriction sites for plasmids pPFMW and pAHW. The coding sequences of *Imp, bru1* and *pAbp* were cloned using *Kpn*I and *Xba*I into plasmid pAWG. A list of primers used to introduce the coding sequences as well as to introduce mutations in the *Mkrn1* coding sequence is appended (Table S6). To analyze the binding of Mkrn1 to *osk*, either the 3’ UTR was cloned downstream of Firefly coding sequence or the complete *osk* gene was cloned in the same vector backbone of pAc5.1B-EGFP (gift from Elisa Izaurralde, Addgene plasmid #21181). For both, restriction sites of *Kpn*I and *Sal*I were used. To ensure the proper usage of the endogenous polyA signal the *osk* 3’ UTR and the *osk* gene included 220 and 248 nucleotides of the downstream sequence, respectively. The primers used for cloning and to introduce mutations are listed in Table S7.

**Table S6.**
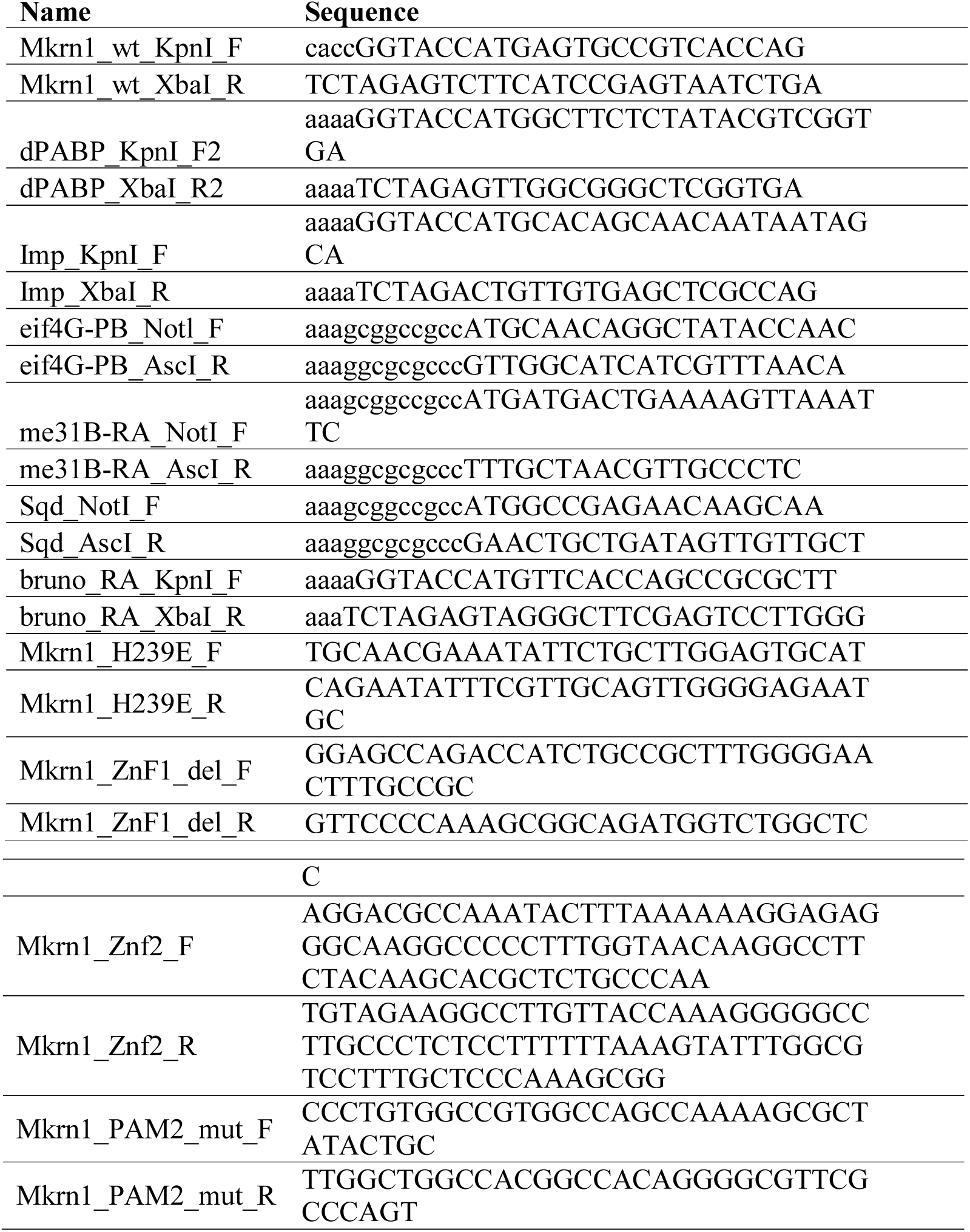
List of primers used to clone CDSs of proteins as well as primers that were used to introduce mutations in the respective CDS.

**Table S7.**
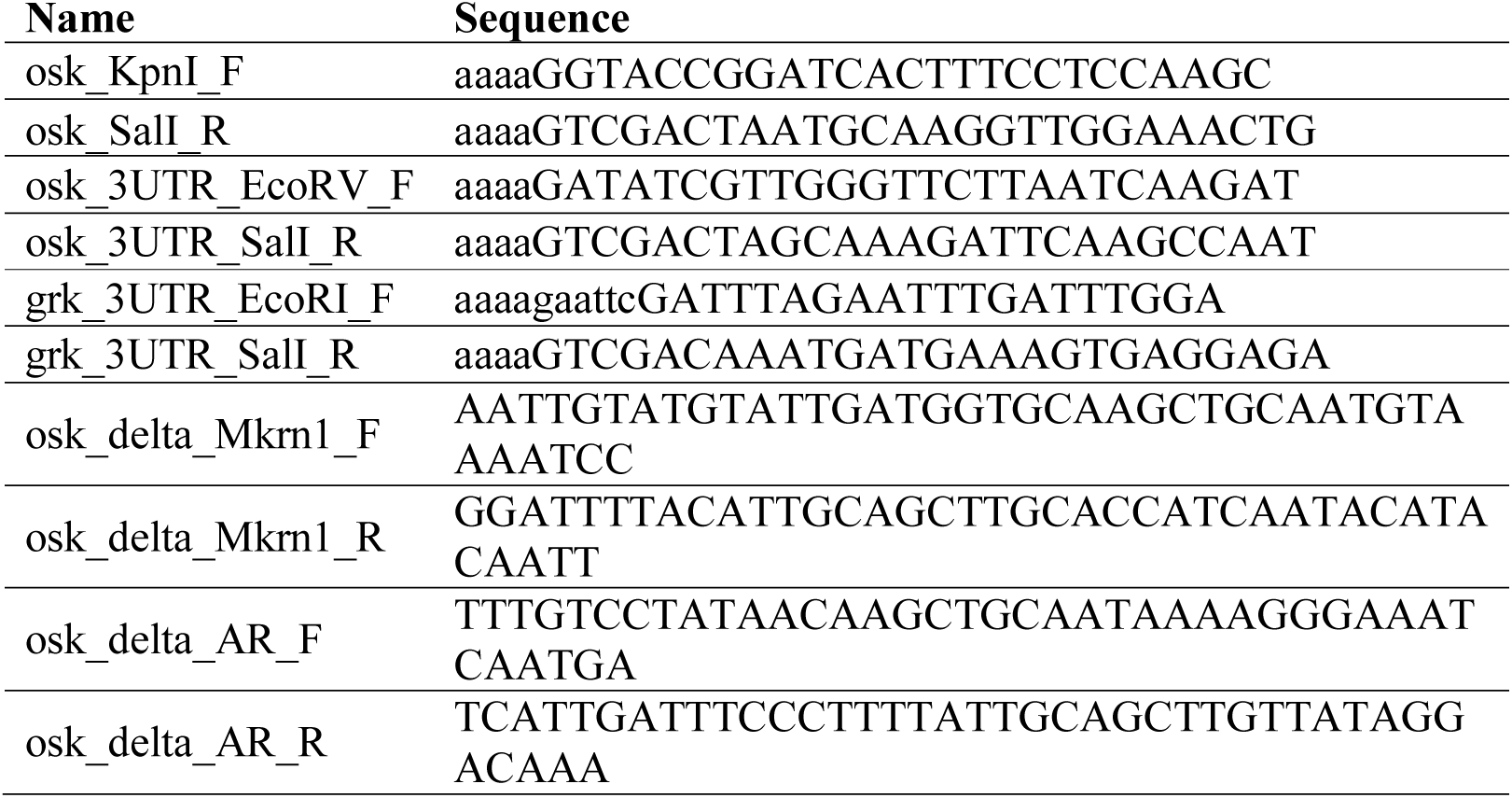
List of primers used to clone *osk* and *grk* reporter genes and introduce the mutations analyzed.

### Acknowledgements

We would like to thank the Bloomington Drosophila Stock Center for fly reagents, and Anne Ephrussi, Craig Smibert, Daniel St Johnston, and Nahum Sonenberg for antibodies. We also thank Violeta Morin and Beili Hu for fly embryo injections, the IMB Proteomics Core Facility for the initial analysis of the mass spectrometry data and the IMB Core Facility Genomics for their helpful support. This work was supported by CIHR grants MOP-44050 and IOP-107945 to P.L., by the Deutsche Forschungsgemeinschaft (DFG) SPP1935 Grants RO 4681/4-1 to J.Y.R., and KO 4566/3-1 to J.K. K.Z. was supported by the LOEWE program Ubiquitin Networks (Ub-Net) of the State of Hesse (Germany) and the SFB 902 of the German Research Foundation.

## Author Contributions

A.D., H.H. and N.L. performed the experiments and wrote the draft of the manuscript.

A.H. performed MS experiments and iCLIP libraries. A.B. analyzed MS experiments.

M.B. and A.B. analyzed iCLIP data. C.R. analyzed and identified the PAM2 motif.

P.B., K.Z. and J.K. guided the MS and iCLIP experiments. J.Y.R. and P.L. supervised the project and finalized the manuscript.

## Supplementary Figure Legends

**Figure S1. Four *Makorin*-related genes in *Drosophila melanogaster***.

(**A**) Sequence alignment of human MKRN1 and Mkrn1, CG5334, CG5347, and CG12477, the four Makorin-related proteins in *Drosophila melanogaster*. The ZnF1 domain in Mkrn1 is highlighted green, the PAM2 motif is highlighted yellow, the RING domain is highlighted red, and the ZnF2 domain is highlighted light blue. The RING and ZnF2 domains are conserved in all four proteins, whereas the PAM2 domain is only conserved in CG12477 and CG5347, and ZnF1 is conserved in CG5334 and CG5347. (**B**) Relative mRNA levels of *Mkrn1* and the three other genes encoding predicted Makorin proteins at various stages of development, as measured by RT-qPCR. mRNA levels were normalized to *Rpl15* mRNA. Error bars depict Stdev, n=3.

**Figure S2. Mkrn1 colocalizes with pole plasm components.**

All images are from wild-type oocytes expressing Venus-Mkrn1 or FLAG-Mkrn1 as indicated. (**A, D, M, P, S, V**) Immunostaining with α-GFP recognizing Venus-Mkrn1. (**G** and **J**) Immunostaining with α -FLAG recognizing FLAG-Mkrn1. (**B** and **E**) Immunostaining with α-Stau. (**H** and **K**) Immunostaining with α-Osk. (**N** and **Q**) Immunostaining with α-Vas. (**T**, and **W**) Immunostaining with α-Aub. (**C, F, I, L, O, R, U, X**) Merged images from two preceding panels.

**Figure S3. Effects of *osk* and *vas* mutations on Mkrn1 localization.**

**(A**) Posterior accumulation of either Venus-Mkrn1 or FLAG-Mkrn1 is normal in *osk*^*54*^*/TM3,Sb* (*osk/+*) oocytes but is absent in *osk*^*54*^*/Df(3R)p-XT103* (*osk*) oocytes. (**B**) Posterior accumulation of either Venus-Mkrn1 or FLAG-Mkrn1 is normal in both *vas*^*1*^*/+* (*vas/+*) *or vas*^*1*^*/vas*^*PH*^ (*vas*) oocytes.

**Figure S4. Mkrn1 genetically interacts with *osk* and *vas*.**

(**A**) Pole cell counts from embryos produced by females with the genotypes listed. Embryos from trans-heterozygotes for *Mkrn1* and *osk* or *vas* mutations have fewer pole cells than those from single heterozygote controls. Depicted is Stdev, n=60. (**B**-**J**) Multiple images of *Mkrn1* mutant egg chambers showing variability of effects on Stau and Grk localization. (**B**-**D**) *Mkrn1*^*W*^ egg chambers immunostained with α-Stau. (**E**-**G**) Multiple images of *Mkrn1*^*W*^ egg chambers immunostained with α-Grk. (**H**-**J**) Multiple images of *Mkrn1*^*S*^ egg chambers immunostained with α-Grk.

**Figure S5. Transgenic expression of tagged Mkrn1 rescues all *Mkrn1* mutant phenotypes**.

(**A**-**C**) Bright-field micrographs of entire ovaries from (**A**) *Mkrn1*^*N*^; (**B**) *nos>FLAG-Mkrn1*; *Mkrn1*^*N*^ (**C**) wild-type females, showing overall rescue of oogenesis. (**D**-**F**) α - Osk immunostaining on (**D**) *Mkrn1*^*W*^, (**E**) *Mkrn1*^*S*^, (**F**) *Mkrn1*^*N*^ egg chambers as negative controls. (**G**-**J**) Transgenic expression of tagged *Mkrn1* restores posterior localization of Osk protein in *Mkrn1*^*W*^ oocytes. (**G, H**) *nos>Venus-Mkrn1; Mkrn1*^*W*^; (**I**, and **J**) *nos>FLAG-Mkrn1; Mkrn1*^*W*^. (**H** and **J**) Immunostaining with α -Osk; (**G**) Immunostaining with α -GFP recognizing Venus-Mkrn1; (**I**) Immunostaining with α - FLAG recognizing FLAG-Mkrn1. (**K**-**N**) Transgenic expression of tagged *Mkrn1* restores expression and posterior localization of Osk protein in *Mkrn1*^*S*^ oocytes. (**K** and **L**) *nos>Venus-Mkrn1*; *Mkrn1*^*S*^; (**M** and **N**) *nos>FLAG-Mkrn1*; *Mkrn1*^*S*^. (**L** and **N**) Immunostaining withα -Osk; (**K**) Immunostaining with α -GFP recognizing Venus-Mkrn1; (**M**) Immunostaining with α -FLAG recognizing FLAG-Mkrn1. (**O** and **P**) Transgenic expression of tagged *Mkrn1* restores expression and posterior localization of Osk protein in *Mkrn1*^*N*^ oocytes. (**Q**-**T**) Immunostaining experiments revealing localization of various proteins in *nos>Venus-Mkrn1; Mkrn1*^*N*^ oocytes. (**Q**) α -Stau; (**R**) α -Vas; (**S**) α -Aub; (**T**) α -Grk.

**Figure S6. *Mkrn1* mutations affect accumulation of mRNAs involved in axis patterning in embryos.**

(**A** and **B**) Anterodorsal accumulation of *grk* is similar to wild-type in stage 10 *Mkrn1*^*W*^ oocytes. Scale bars, 50 μm. (**C**) *grk mRNA* remains associated with the oocyte nucleus and is mislocalized to the posterior in stage 10 *Mkrn1*^*S*^ oocytes. Scale bars, 50 μm. *In situ* hybridization experiments showing posterior accumulation of (**D**) *osk*, (**E**) *nos*, and (**F**) *pgc* mRNAs in wild-type embryos. (**G-I**) Posterior accumulation of these mRNAs is lost in *Mkrn1*^*W*^ embryos.

**Figure S7. Interactome of Mkrn1 in S2R+ cells.**

(**A**) Schematic diagram of Mkrn1 constructs with functional domains highlighted. Mkrn1^RING^ carries a point mutation that changes histidine 239 to glutamic acid (H239E) while Mkrn1ΔZnF1 contains a deletion of amino acids 26 to 33. To disrupt the ZnF2 domain (Mkrn1^ZnF2^) three point mutations that change the cysteines to alanines at positions 302, 312 and 318 (C302A, C312A and C318A) were introduced. (**B**) Immunoblot showing the relative expression levels of various forms of FLAG-Mkrn1 in S2R+ cells. (**C, D**) Volcano plots showing the interactome of (**C**) Myc-Mkrn1^ING^ and (**D**) Myc-Mkrn1 in S2R+ cells as identified using mass spectrometry and label-free quantification. The enrichment of proteins compared to control was plotted in a volcano plot using a combined cutoff of log_2_ fold change of 2 and an FDR of 0.05. Several proteins of interest are labelled. The entire list of enriched proteins can be found in Supplementary Tables 1 and 2.

**Figure S8. Validation of Mkrn1 interactome.**

Pulldown experiments to validate associations of tagged Mkrn1^RING^ with (**A**) GFP-pAbp, (B) GFP-Imp, (**C**) Myc-eIF4G (**D**) Myc-Sqd and (**E**) Myc-Me31B. GFP and Myc immunoprecipitation was performed in the absence or presence of RNase T1 and enrichment of the proteins was analyzed by immunoblotting. (**F**) Western blot analysis of co-IP experiments between Venus-Mkrn1 and eIF4G. α-Tubulin (αtub, lanes 1, 2) and ovaries lacking the *Venus-Mkrn1* transgene (lane 4) were used as negative controls.

**Figure S9. Analysis of interaction strength between Mkrn1**^**RING**^ **and binding partners.**

Representative western blots of co-IP experiments between Mkrn1^RING^ and (**A**) Myc-pAbp, (**B**) Myc-eIF4G, (**C**) Myc-me31B, (**D**) Myc-Sqd and (E) GFP-Imp using either 150 mM, 300 mM or 500 mM salt concentration for washing. The intensity of the Mkrn1^RING^ signal after IP was quantified and normalized to input levels. The resulting enrichment over control IP is summarized in Figure 5B.

**Figure S10. Analysis of the RNA binding ability of Mkrn1.**

**(A)** The RNA binding activity of Mkrn1 is mediated by its ZnF1 domain. Autoradiographs showing RNA binding to various forms of Mkrn1 and a GFP negative control. Crosslinked RNA-protein complexes were immunoprecipitated with α-FLAG and treated with different dilutions of RNase I (left: 1/50, right: 1/5000). RNA was subsequently radiolabelled and the RNA-protein complexes were separated by SDS-PAGE. Bound RNA of different sizes is detected by a smear extending upward from the sharp bands that correspond to the sizes of the FLAG-Mkrn1 proteins (arrow). (**B**) Representative immunoblot of RIP experiments shown in Figure 6A. Either Mkrn1 or Mkrn1^ΔZnF1^ were overexpressed in *Mkrn1*^*N*^ ovaries using *nos>Gal4* driver. The proteins were pulled down using α-FLAG antibody. The deletion protein is larger because of the presence of the additional Myc tag. (**C**) Validation of iCLIP experiments. Immunoprecipitation of FLAG-Mkrn1 was performed in different conditions. S2R+ cells were transfected and UV-crosslinked prior to IP experiments. Left: autoradiograph showing protein-RNA complexes. Right: Signals of lanes 2, 4 and 5 in autoradiograph were cut and RNA isolated. RNA length was analyzed on a TBE-urea gel. (**D**) iCLIP datasets with Mkrn1-FLAG in S2R+ cells. The x axis displays maximum binding strength per gene (SOB) and the y axis shows gene identity. The genes are sorted by SOB with *osk* mRNA appearing at the third place. Note that ribosomal genes have been excluded for clarity.

**Figure S11. Mkrn1 binding to *osk* 3’ UTR is dependent on pAbp.**

(**A**) Mkrn1 but not Mkrn1^ΔZnF1^ binds to the *osk* 3’ UTR in S2R+ cells. FLAG-RIP of GFP, Mkrn1 and Mkrn1^ΔZnF1^ was performed in S2R+ cells co-expressing *luciferase*-*grk*-3’UTR or *luciferase*-*osk*-3’UTR reporter. Error bars depict Stdev, n=3. (**B**-**E**) Western blot analysis of a representative RIP experiment summarized in Figures 6C-E. RIP of GFP or Mkrn1^RING^ were performed either in presence of *luciferase-osk*-3’ UTR with wild-type sequence, (**B**) a mutation in the Mkrn1 binding site (*osk*^ΔMkrn1^) or (**C**) a mutation in the AR region (*osk*^ΔAR^). RIP experiments against FLAG-Mkrn1^RING^ were performed in control (*LacZ*) condition and compared to (**D**) *Imp* or (**E**) *pAbp* mRNA knockdown. Right: RT-qPCR analysis of the knockdown efficiency. *Imp* and *pAbp* mRNA levels were normalized to *rpl15* mRNA. (**F**) FLAG-RIP experiments in S2R+ cells using different *Mkrn1* mutants. Representative immunoblot is depicted.

**Figure S12. Binding of Bru1 to *osk* 3’UTR is antagonized by Mkrn1.**

(**A**) RIP experiment of GFP alone (ctrl) or GFP-tagged Bru1 in S2R+ cells. Left: qPCR analysis of RIP experiment analyzing co-transfected *luciferase-osk*-3’UTR and endogenous *grk* mRNA. Error bars depict Stdev, n=3. Right: Immunoblot of the IP. (**B**-**D**) Immunoblots of representative RIP experiments summarized in Figures 7A-C. Right: RT-qPCR validation of the respective knockdown. mRNA levels were normalized to *rpl15* mRNA. (**B**) RIP experiment of either GFP alone, GFP-Imp or GFP-Bru1 was performed in control (*LacZ*) or *Mkrn1* mRNA knockdown condition. (**C**) RIP experiment performed against FLAG-tagged pAbp or Sqd in knockdown condition of *LacZ* or *Mkrn1* mRNA. (**D**) GFP-RIP against either GFP alone or GFP-Bru1 in control or *pAbp* KD. (**E**) Representative immunoblot of a RIP experiment using control or *Mkrn1*^*W*^ ovary lysate against endogenous Bru1. As control IP, rabbit IgG was used. (**F-H**) The three panels show the same *Mkrn1*^*W*^ stage 10 egg chamber stained for (**F**) Stau, (**G**) *osk* mRNA and (**H**) a merged image. There is accumulation of Stau near the pole plasm and co-localization with *osk* mRNA.

## Supplementary Tables

**Supplementary Table 1.** List of identified interaction partners of Mkrn1^RING^ using mass spectrometry and limma analysis. All proteins with an FDR ≤ 0.05 05 and log_2_ fold change ≥ 2 are depicted. (see supplementary file 1)

**Supplementary Table 2.** List of identified interaction partners of Mkrn1 using mass spectrometry and limma analysis. All proteins with an FDR ≤ 0.05 and log_2_ fold change ≥ 2 are depicted. (see supplementary file 1)

