## Supplementary Figures 1-12 for "Makorin 1 controls embryonic patterning by alleviating Bruno1-mediated repression of *oskar* translation"

Dold_Fig_S1

**A**

```
Hs MKRN1          MAEAATPGTTATTSGAGAAAATAAAASPTPIPTVTAPSLGAGGGGGGSDGSGGGWTKQVT    60
Dm CG12477        ------------------------------------------------------------    0
Dm Mkrn1          ------------------------------MSAVTSSDTAI--------SGMALGRSQTI    22
Dm CG5334         -------------------------------------------------MTSSRQNLHKI    11
Dm CG5347         ---------------------------------------------------MSYRRSQTI    9

Hs MKRN1          CRYFMHGVCKEGDNCRYSHDLSDSPYSVVCKYFQRGYCIYGDRCRYEHSKPLKQEEATAT    120
Dm CG12477        --------------------MS----TVASN-----------------------DQHVDT    13
Dm Mkrn1          CRYYVRGICRFGELCRFSHDLSRGRPECEEQ-----------------------VAT---    56
Dm CG5334          CRYRLRGNCRFDDLCRYSHDVL-QNDNLRPQ-----------------------ISNNAD    47
Dm CG5347          CRFHLLGICRFGDLCRFSHDETTPNDNQSPQ-----------------------ISEIAD    46
                                                ：

Hs MKRN1          ELTTKSSLAASSSLSSIVGPLVEMNTGEAESRNSNFATVGAGSEDWVNAIEFVPGQPYCG    180
Dm CG12477        GVLPVPS----------------------------TNSPSSQIEQKGNEAIVPNYKAAT    44
Dm Mkrn1          DVLPKPSTSSSS------------------TIGSRSASISSQQRNWANAPVFVPSQKRYT    98
Dm CG5334          EVVGHA------------------------------NRYSRPMTGA------------    63
Dm CG5347          EVVENE------------------------QVVASTSSYSRQMTWANAPEFVPRYKANS    81
                   :                                    .  :

Hs MKRN1          RTAPSCTEAP--------------------LQGSVTKEESEKEQTAVETKKQLCPYAAVG    220
Dm CG12477        AGEQGEAQAG------------ATCTTVAVRPSGVATSADIVKGSSS-------------    79
Dm Mkrn1          AHEQSEFETTVDPEAVMEAQAGASYDTLAP---GV-SWAEVVGGPSSLNKEDYGEE--NS    152
Dm CG5334          ------------------------------------------------------------    63
Dm CG5347          ADFQA----T------------------------------------EGVQDICPY--GG    98

Hs MKRN1          ECRYGENCVY-LHGDSCDMCGLQVLHPMDAAQRSQHIKSCIEAHEKDMELSFAVQRSKDM    279
Dm CG12477        ----------------------------------------VSSH-VQPSWRSSFARSQDK    98
Dm Mkrn1          SCAWGEFSAYPIHMELCEMCDQYCLHPTDQVQRRSHNRECLQQHEQAMELSFAIARSKDK    212
Dm CG5334          ------------------------------------------NATEEMKLSFAIAKSQDK    81
Dm CG5347          SCIWGSKCSYPLHMEICKMCDLYCLHPMDQNQRRAHNRECLEQHEQAMELSFAIARSKDK    158
                                                                 .   :. :*:*

Hs MKRN1          VCGICMEVVYEKANPSERRFGILSNCNHTYCLKCIRKWRSAKQFESKIIKSCPECRITSN    339
Dm CG12477        KCGICFETIMEKEG-GDKRFGILPSCNHVFCFQCICTWRHATQYAYQVTRACPECRVWSN    157
Dm Mkrn1          TCGICFDTIMEKAG-REKRFGILPNCNHIFCLECIRTWRQAKQFENKITRACPECRVCSD    271
Dm CG5334          MCGICLETVVKKRG-RECRFGILPKCKHIFCLTCIRRWRQAEYIEDNVKRGCPECRVFSE    140
Dm CG5347          MCGICFDTVVEKKG-RERRFGILSKCKHIFCLTCIRTWRQAHQFEATVTRGCPECRVFSE    217
                    ****::.: :* .  : ***** .*:* :*: **  ** *      : :.*****: *:

Hs MKRN1          FVIPSEYWVEEKEEKQKLILKYKEAMSNKACRYFDEGRGSCPFGGNCFYKHAYPDGRREE    399
Dm CG12477        FVCPSAFWVEEKVAKDQLINDHLAAMRARDCKYFKQGQGVCLFGNKCFYKHSIPNADYVD    217
Dm Mkrn1          FVCPSAFWMETKEEKDKLLNDYRAALGAKDCKYFKKGEGKCPFGNKCFYKHALPNGDIVD    331
Dm CG5334          FVCPSAYWVDTKEEKDKLLSEYRAAMGAKDCKYFNGGLGKCPFGTNCFYNHRSLAGDNRI    200
Dm CG5347          FVCPSAYWVDTKEEKDKLLSEYRAAMGAKDCKYFNGGLGKCPFGNKCFYRHHRRCDDNQT    277
                   ** ** :*:: *  *::*: .:  *:  : *:**. * * * ** :***.*

Hs MKRN1          PQRQKVGTS----SR---------YRAQRRNHFWELIEERENSNPFDNDEEEVVTFELGE    446
Dm CG12477        VGLPTHALGLPIPSDFSGLGNCLILVPFPNMFFDDFSNSDDY--------------DFSD    263
Dm Mkrn1          VGLPKRTRKLQSQNEIID---------LLDIYLWDYVDRRDY--------------HWLE    368
Dm CG5334          QRHSMEDDDF--------------------------------------------------    210
Dm CG5347          QSNAVNSD----------------------------------------------------    285

Hs MKRN1          MLLMLLAAGGDDELTDSEDEWDLFHDELEDFYDLDL    482
Dm CG12477        VN----------------------------------    265
Dm Mkrn1          MISSDITS-----------------SESSDYSDED-    386
Dm CG5334          ------------------------------------    210
Dm CG5347          ------------------------------------    285
```

**B**

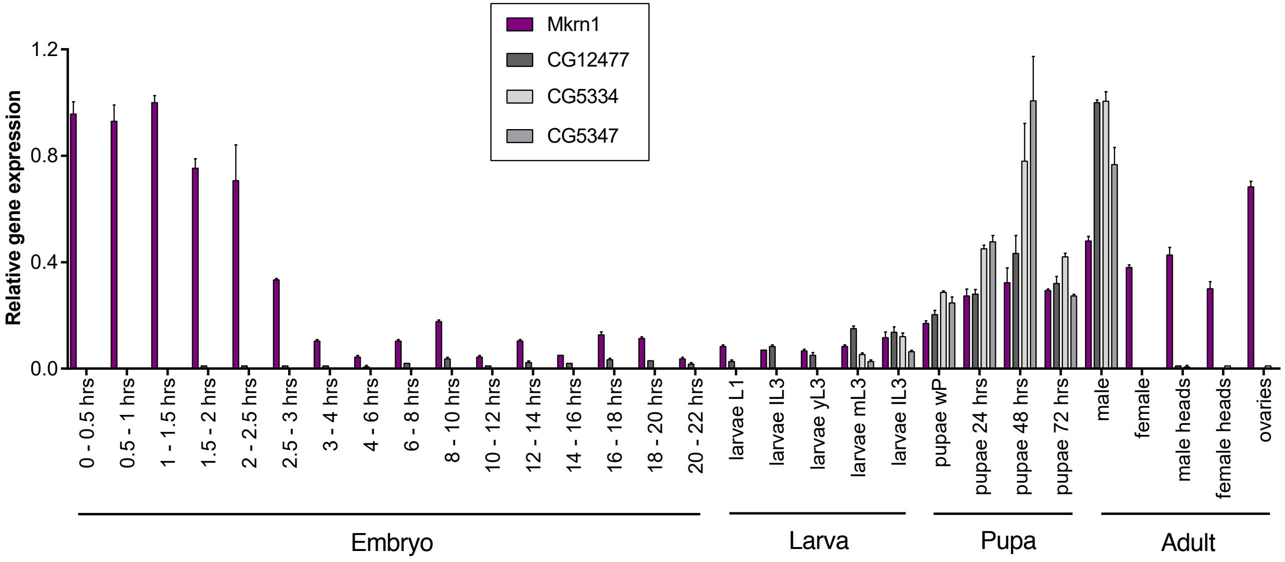

Dold_Fig_S2

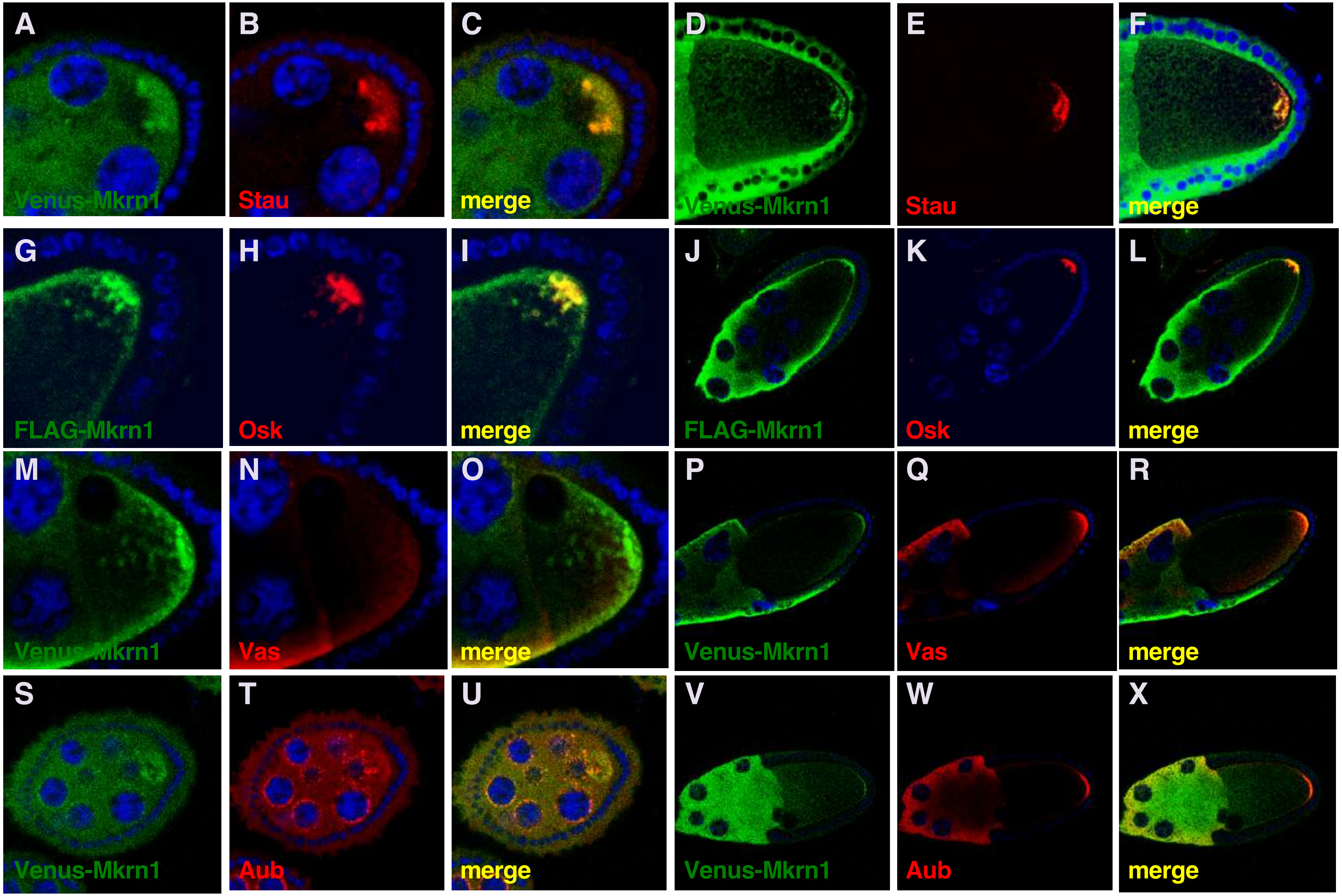

A
Venus-Mkrn1
B
Stau
C
merge
D
Venus-Mkrn1
E
Stau
F
merge
G
FLAG-Mkrn1
H
Osk
I
merge
J
FLAG-Mkrn1
K
Osk
L
merge
M
Venus-Mkrn1
N
Vas
O
merge
P
Venus-Mkrn1
Q
Vas
R
merge
S
Venus-Mkrn1
T
Aub
U
merge
V
Venus-Mkrn1
W
Aub
X
merge

Dold_Fig_S3

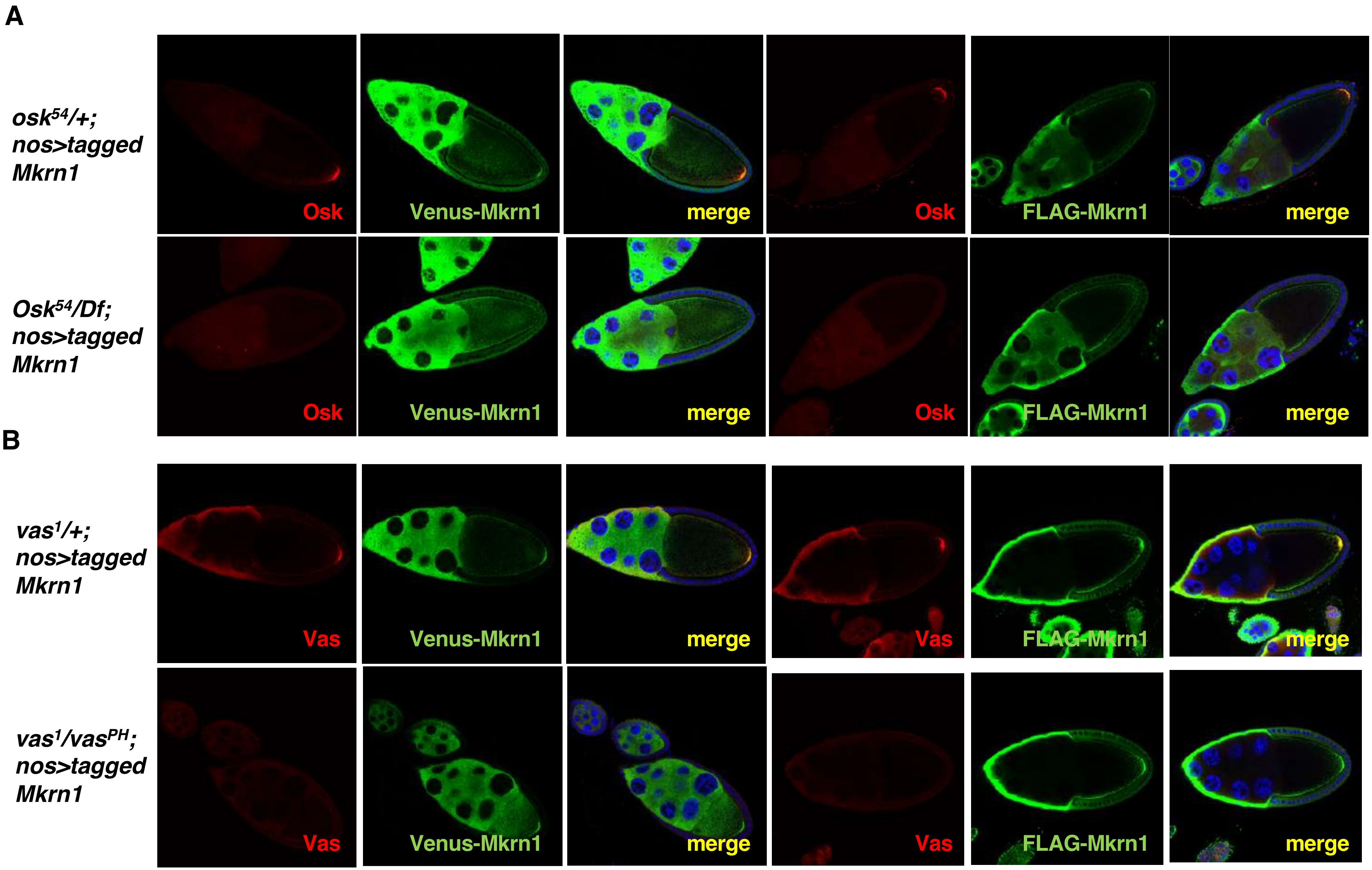
A
osk54/+;
nos>tagged
Mkrn1
Osk
Venus-Mkrn1
merge
Osk
FLAG-Mkrn1
merge
Osk54/Df;
nos>tagged
Mkrn1
Osk
Venus-Mkrn1
merge
Osk
FLAG-Mkrn1
merge
B
vas1/+;
nos>tagged
Mkrn1
Vas
Venus-Mkrn1
merge
Vas
FLAG-Mkrn1
merge
vas1/vasPH;
nos>tagged
Mkrn1
Vas
Venus-Mkrn1
merge
Vas
FLAG-Mkrn1
merge

Dold_Fig_S4

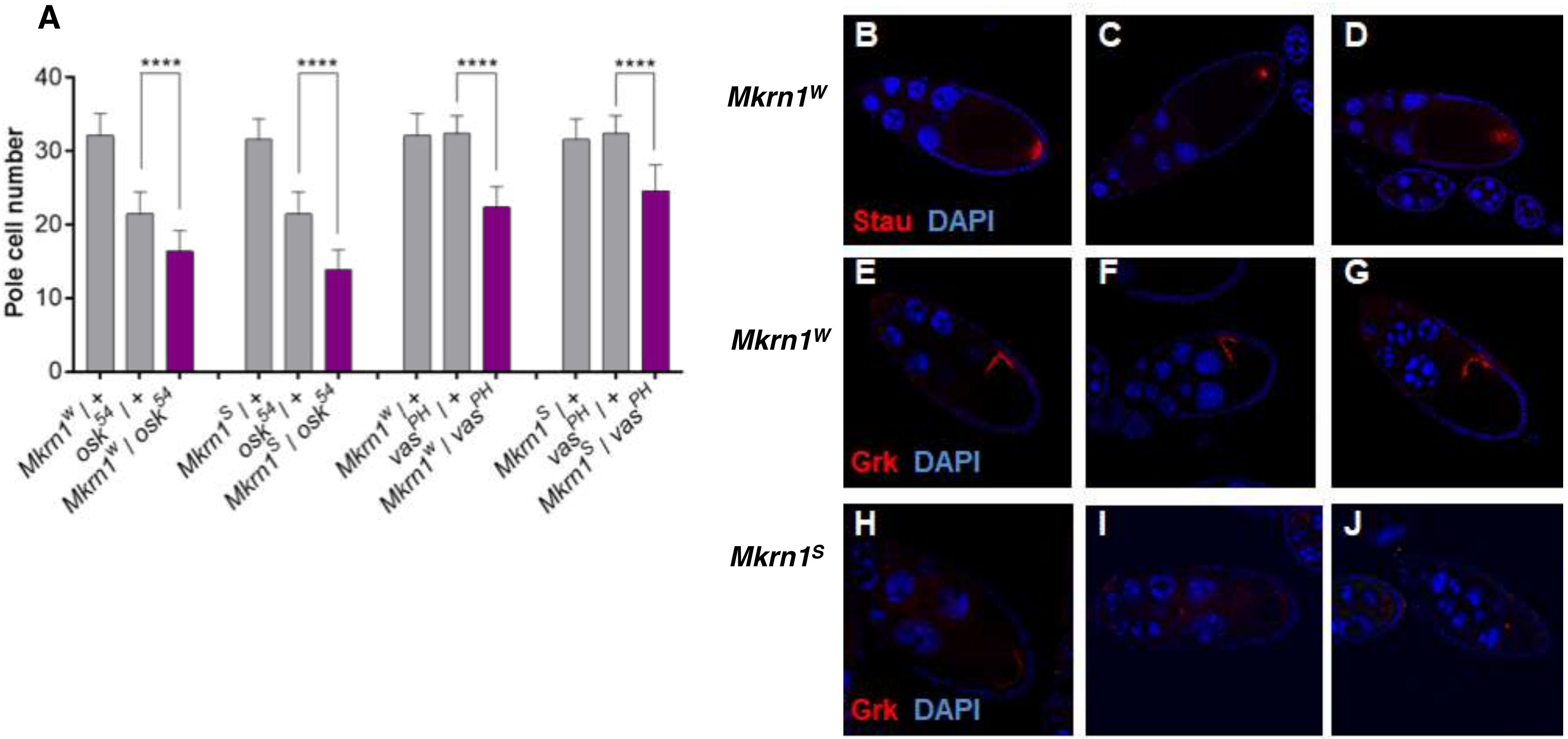
A
****
****
****
****
Pole cell number
40
30
20
10
0
Mkrn1^W / +
osk^54 / +
Mkrn1^W / osk^54
Mkrn1^S / +
osk^54 / +
Mkrn1^S / osk^54
Mkrn1^W / +
vas^PH / +
Mkrn1^W / vas^PH
Mkrn1^S / +
vas^PH / +
Mkrn1^S / vas^PH
Mkrn1^W
Mkrn1^W
Mkrn1^S
B
C
D
Stau DAPI
E
F
G
Grk DAPI
H
I
J
Grk DAPI

Dold_Fig_S5

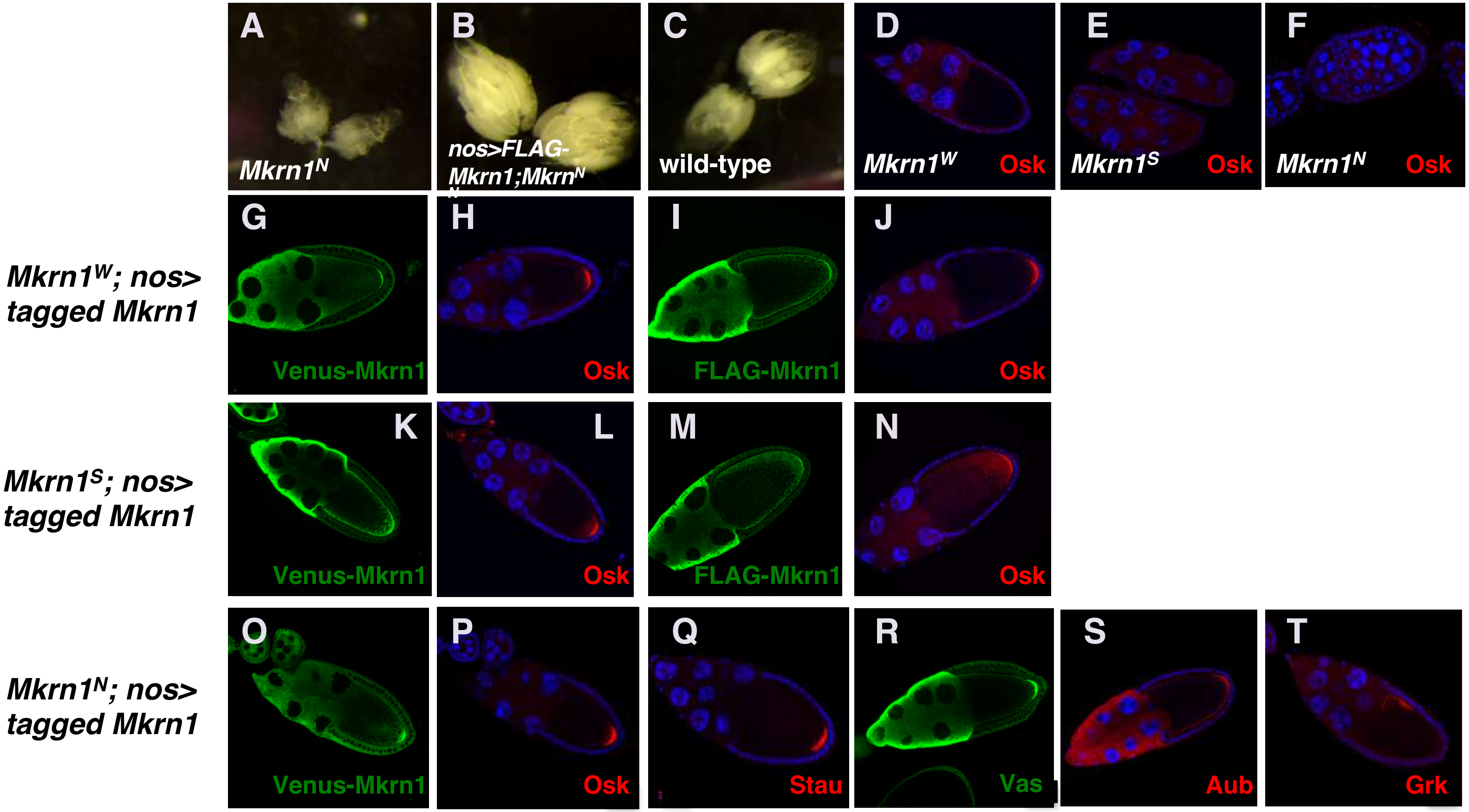
A
Mkrn1N
B
nos>FLAG-Mkrn1;MkrnN
C
wild-type
D
Mkrn1W
Osk
E
Mkrn1S
Osk
F
Mkrn1N
Osk
Mkrn1W; nos> tagged Mkrn1
G
Venus-Mkrn1
H
Osk
I
FLAG-Mkrn1
J
Osk
Mkrn1S; nos> tagged Mkrn1
K
Venus-Mkrn1
L
Osk
M
FLAG-Mkrn1
N
Osk
Mkrn1N; nos> tagged Mkrn1
O
Venus-Mkrn1
P
Osk
Q
Stau
R
Vas
S
Aub
T
Grk

Dold_Fig_S6

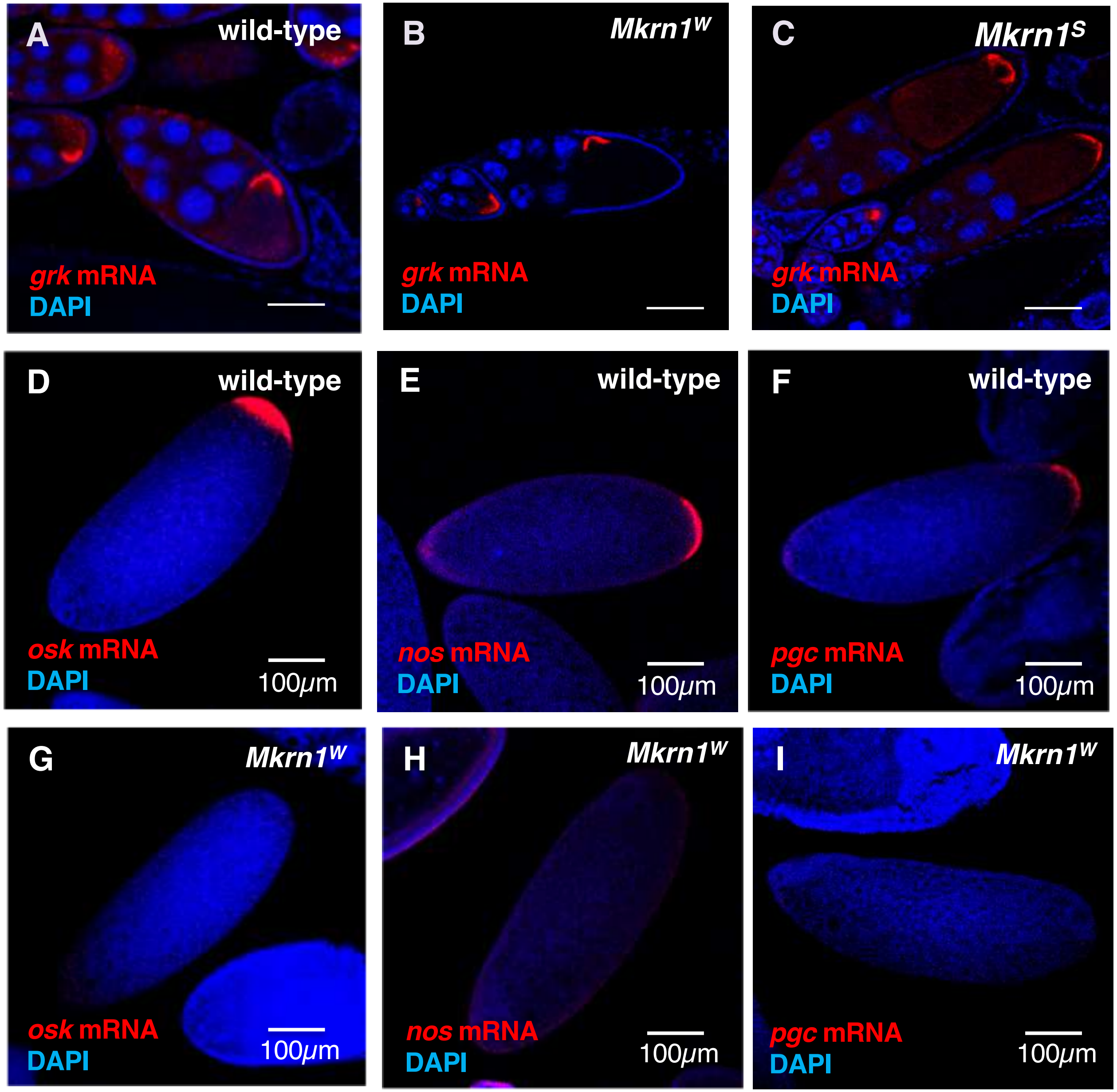

A
wild-type
grk mRNA
DAPI
B
Mkrn1W
grk mRNA
DAPI
C
Mkrn1S
grk mRNA
DAPI
D
wild-type
osk mRNA
DAPI
100μm
E
wild-type
nos mRNA
DAPI
100μm
F
wild-type
pgc mRNA
DAPI
100μm
G
Mkrn1W
osk mRNA
DAPI
100μm
H
Mkrn1W
nos mRNA
DAPI
100μm
I
Mkrn1W
pgc mRNA
DAPI
100μm

Dold_Fig_S7

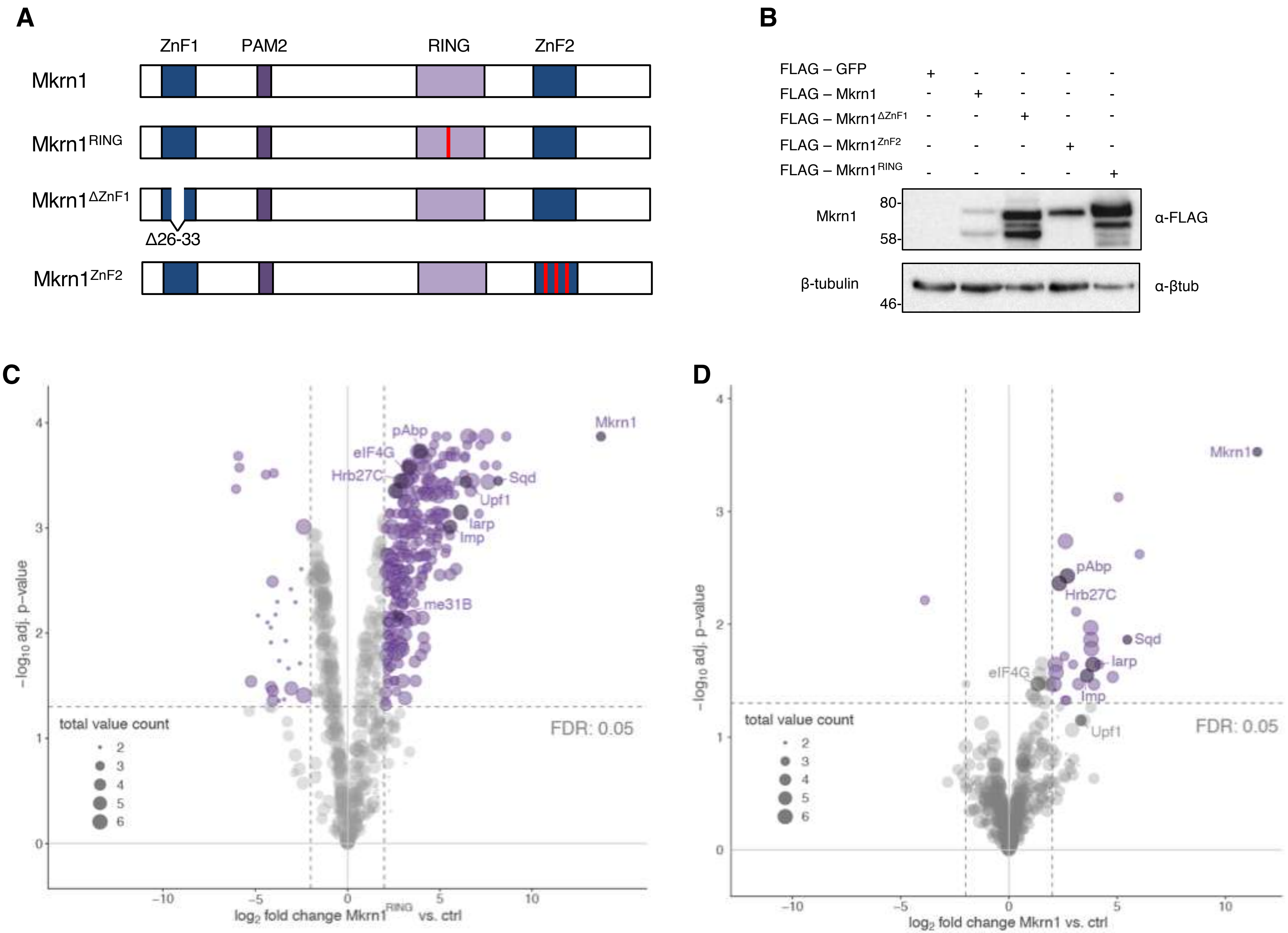
A
ZnF1
PAM2
RING
ZnF2
Mkrn1
Mkrn1RING
Mkrn1ΔZnF1
Δ26-33
Mkrn1ZnF2
B
FLAG – GFP
FLAG – Mkrn1
FLAG – Mkrn1ΔZnF1
FLAG – Mkrn1ZnF2
FLAG – Mkrn1RING
Mkrn1
80-
58-
α-FLAG
β-tubulin
46-
α-βtub
C
Mkrn1
pAbp
eIF4G
Hrb27C
Sqd
Upf1
larp
Imp
me31B
total value count
FDR: 0.05
−log10 adj. p−value
log2 fold change Mkrn1RING vs. ctrl
D
Mkrn1
pAbp
Hrb27C
Sqd
larp
eIF4G
Imp
Upf1
total value count
FDR: 0.05
−log10 adj. p−value
log2 fold change Mkrn1 vs. ctrl

Dold_Fig_S8

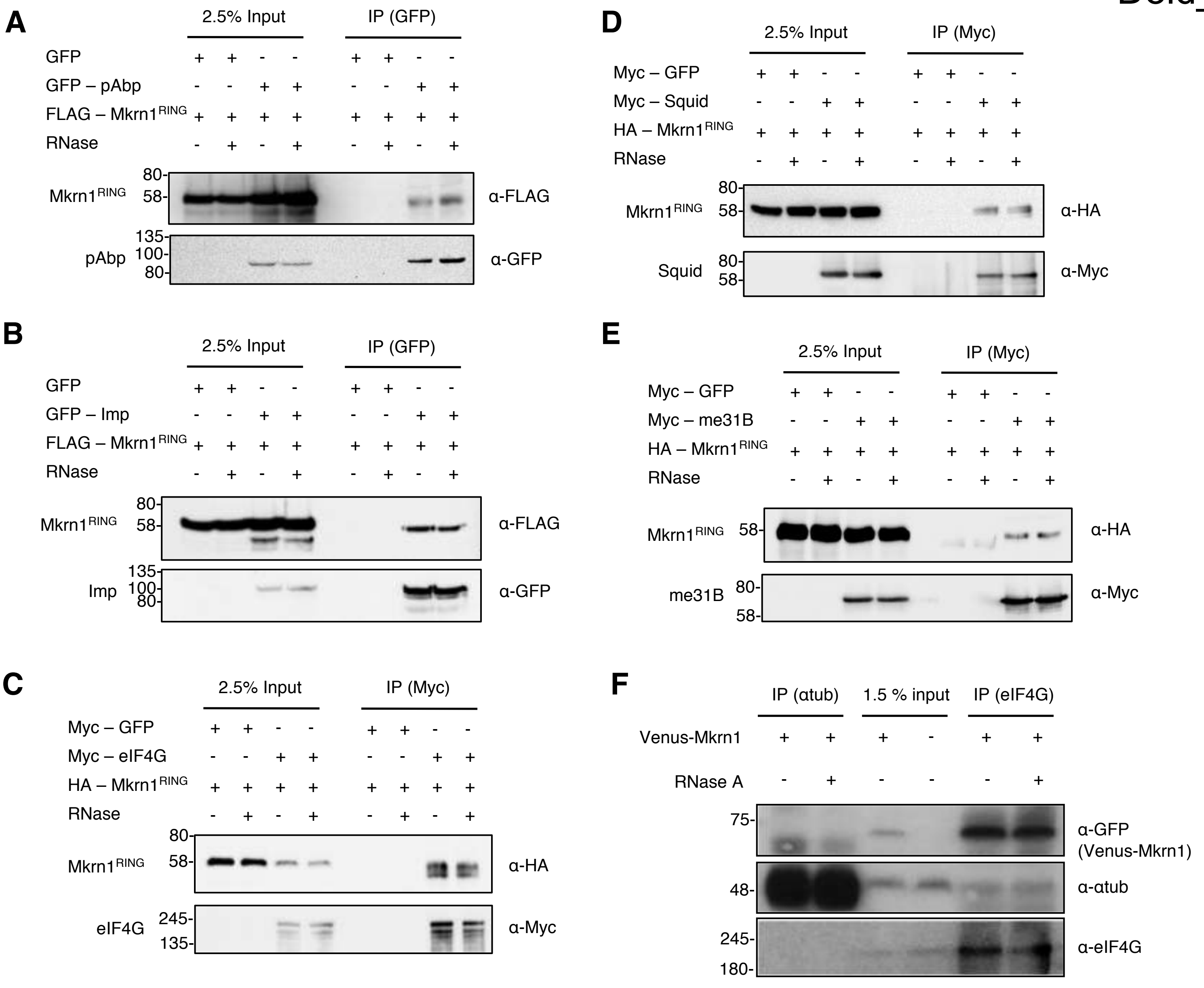
A
2.5% Input
IP (GFP)
GFP
GFP – pAbp
FLAG – Mkrn1RING
RNase
Mkrn1RING
pAbp
80-
58-
135-
100-
80-
α-FLAG
α-GFP
B
2.5% Input
IP (GFP)
GFP
GFP – Imp
FLAG – Mkrn1RING
RNase
Mkrn1RING
Imp
80-
58-
135-
100-
80-
α-FLAG
α-GFP
C
2.5% Input
IP (Myc)
Myc – GFP
Myc – eIF4G
HA – Mkrn1RING
RNase
Mkrn1RING
eIF4G
80-
58-
245-
135-
α-HA
α-Myc
D
2.5% Input
IP (Myc)
Myc – GFP
Myc – Squid
HA – Mkrn1RING
RNase
Mkrn1RING
Squid
80-
58-
80-
58-
α-HA
α-Myc
E
2.5% Input
IP (Myc)
Myc – GFP
Myc – me31B
HA – Mkrn1RING
RNase
Mkrn1RING
me31B
58-
80-
58-
α-HA
α-Myc
F
IP (αtub)
1.5 % input
IP (eIF4G)
Venus-Mkrn1
RNase A
75-
48-
245-
180-
α-GFP
(Venus-Mkrn1)
α-αtub
α-eIF4G

Dold_Fig_S9

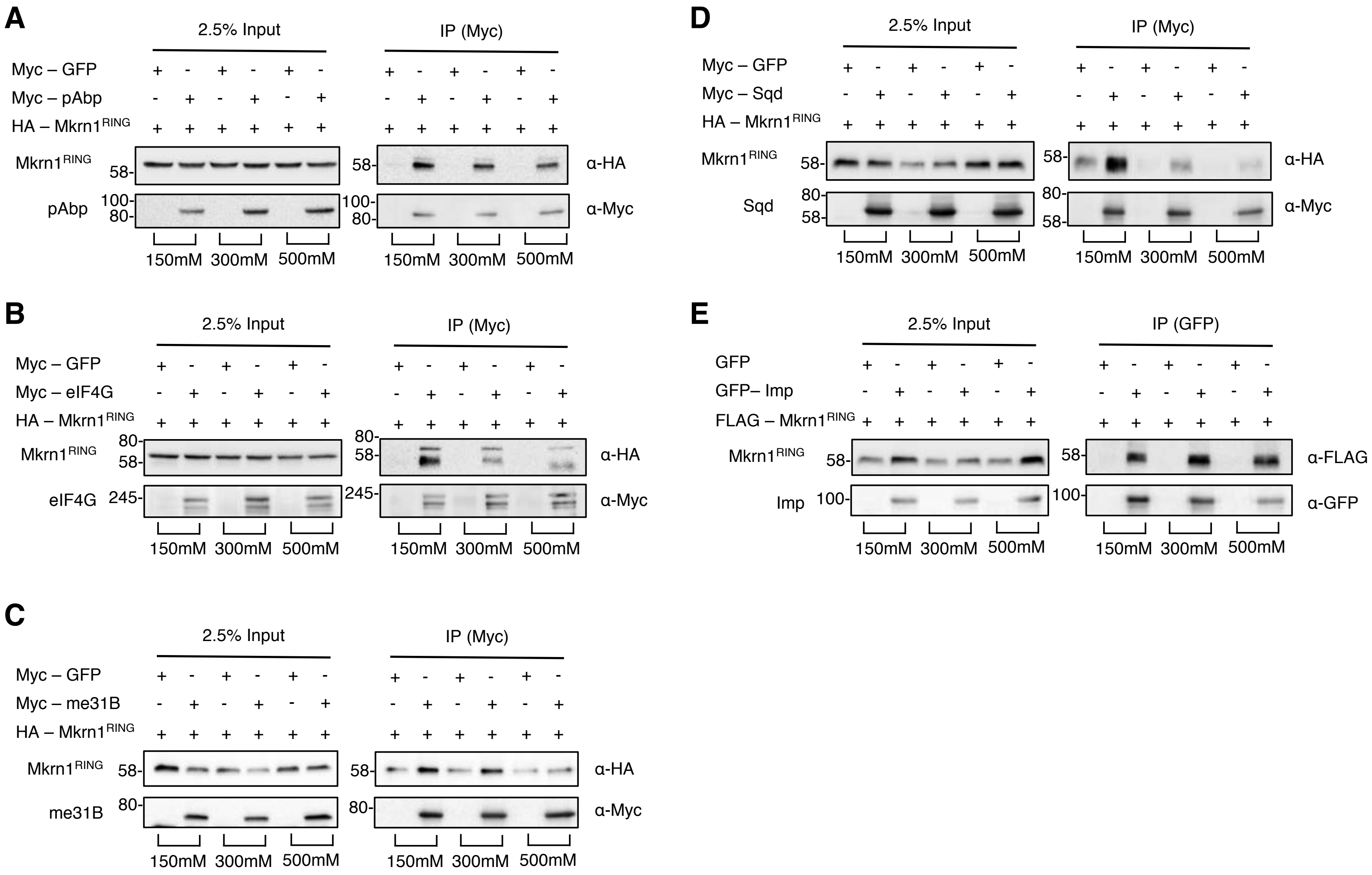

A
2.5% Input
IP (Myc)
Myc – GFP
Myc – pAbp
HA – Mkrn1RING
Mkrn1RING
pAbp
α-HA
α-Myc
150mM 300mM 500mM
B
2.5% Input
IP (Myc)
Myc – GFP
Myc – eIF4G
HA – Mkrn1RING
Mkrn1RING
eIF4G
α-HA
α-Myc
150mM 300mM 500mM
C
2.5% Input
IP (Myc)
Myc – GFP
Myc – me31B
HA – Mkrn1RING
Mkrn1RING
me31B
α-HA
α-Myc
150mM 300mM 500mM
D
2.5% Input
IP (Myc)
Myc – GFP
Myc – Sqd
HA – Mkrn1RING
Mkrn1RING
Sqd
α-HA
α-Myc
150mM 300mM 500mM
E
2.5% Input
IP (GFP)
GFP
GFP– Imp
FLAG – Mkrn1RING
Mkrn1RING
Imp
α-FLAG
α-GFP
150mM 300mM 500mM

Dold_Fig_S10

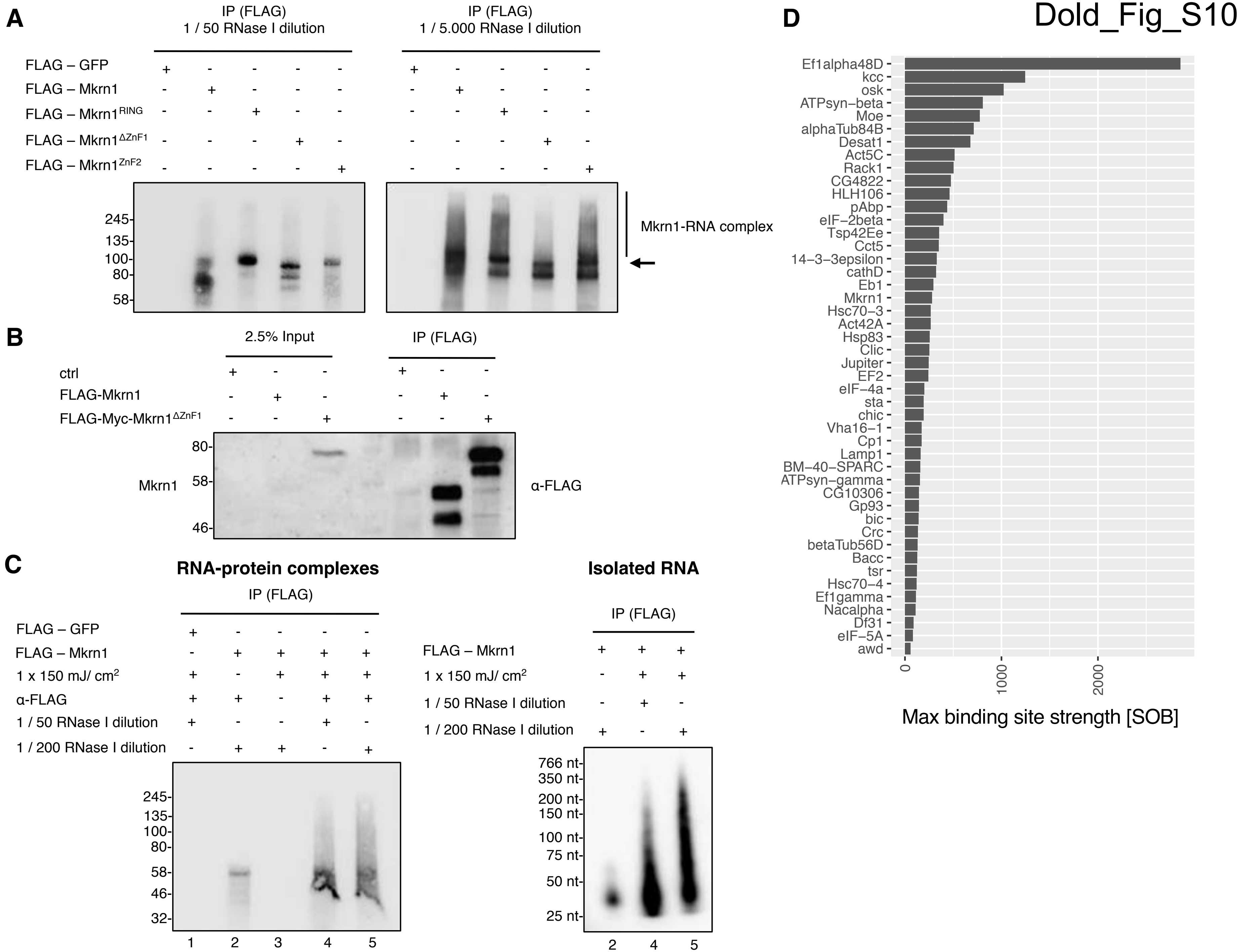
A
IP (FLAG)
1 / 50 RNase I dilution
IP (FLAG)
1 / 5.000 RNase I dilution
FLAG – GFP
FLAG – Mkrn1
FLAG – Mkrn1RING
FLAG – Mkrn1ΔZnF1
FLAG – Mkrn1ZnF2
245
135
100
80
58
Mkrn1-RNA complex
B
2.5% Input
IP (FLAG)
ctrl
FLAG-Mkrn1
FLAG-Myc-Mkrn1ΔZnF1
80
58
46
Mkrn1
α-FLAG
C
RNA-protein complexes
IP (FLAG)
FLAG – GFP
FLAG – Mkrn1
1 x 150 mJ/ cm2
α-FLAG
1 / 50 RNase I dilution
1 / 200 RNase I dilution
245
135
100
80
58
46
32
1 2 3 4 5
Isolated RNA
IP (FLAG)
FLAG – Mkrn1
1 x 150 mJ/ cm2
1 / 50 RNase I dilution
1 / 200 RNase I dilution
766 nt
350 nt
200 nt
150 nt
100 nt
75 nt
50 nt
25 nt
2 4 5
D
Ef1alpha48D
kcc
osk
ATPsyn−beta
Moe
alphaTub84B
Desat1
Act5C
Rack1
CG4822
HLH106
pAbp
eIF−2beta
Tsp42Ee
Cct5
14−3−3epsilon
cathD
Eb1
Mkrn1
Hsc70−3
Act42A
Hsp83
Clic
Jupiter
EF2
eIF−4a
sta
chic
Vha16−1
Cp1
Lamp1
BM−40−SPARC
ATPsyn−gamma
CG10306
Gp93
bic
Crc
betaTub56D
Bacc
tsr
Hsc70−4
Ef1gamma
Nacalpha
Df31
eIF−5A
awd
0
1000
2000
Max binding site strength [SOB]

Dold_Fig_S11

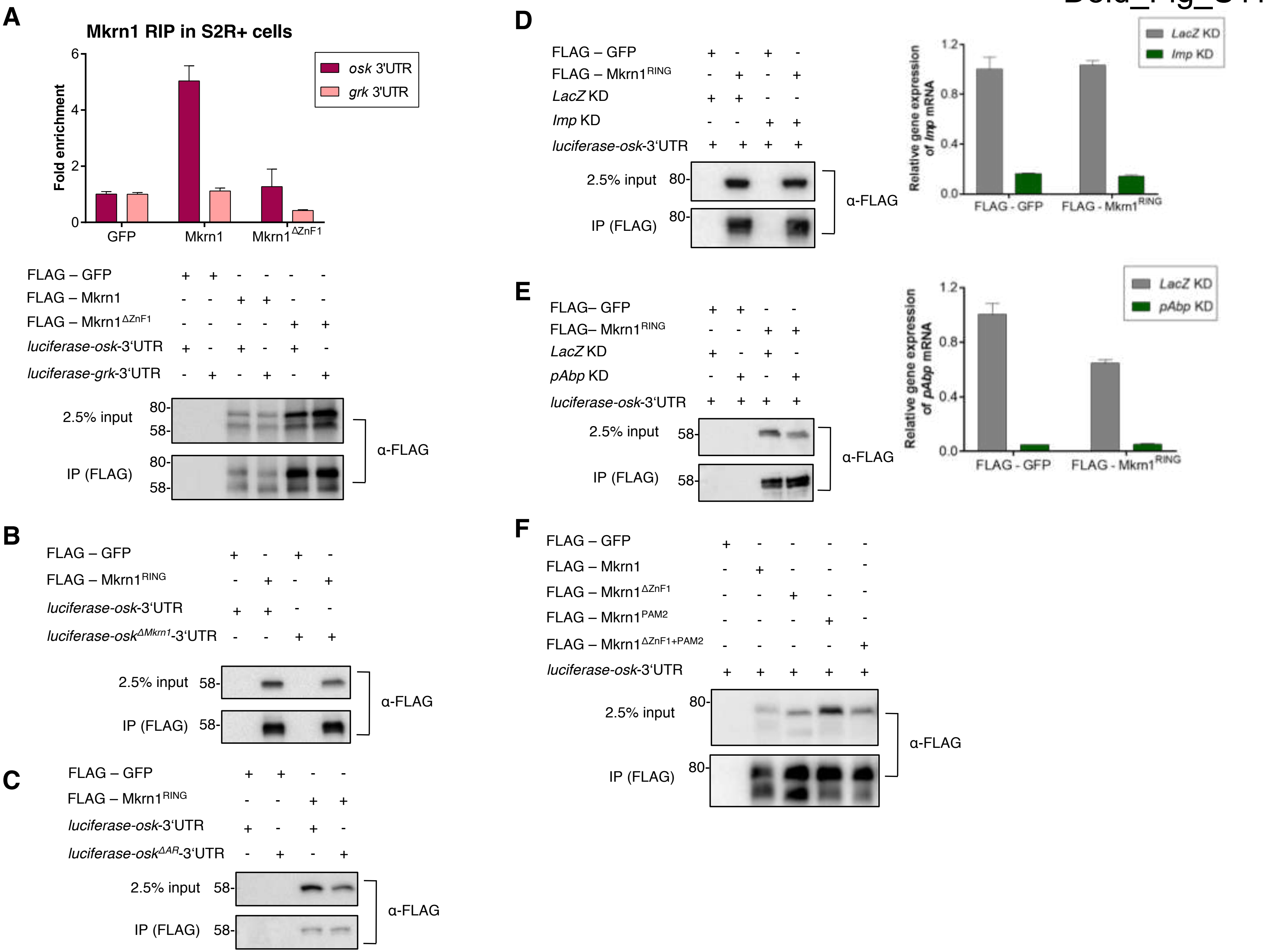
A
Mkrn1 RIP in S2R+ cells
Fold enrichment
osk 3'UTR
grk 3'UTR
GFP
Mkrn1
Mkrn1ΔZnF1
FLAG – GFP
FLAG – Mkrn1
FLAG – Mkrn1ΔZnF1
luciferase-osk-3'UTR
luciferase-grk-3'UTR
2.5% input
IP (FLAG)
α-FLAG
B
FLAG – GFP
FLAG – Mkrn1RING
luciferase-osk-3'UTR
luciferase-oskΔMkrn1-3'UTR
2.5% input
IP (FLAG)
α-FLAG
C
FLAG – GFP
FLAG – Mkrn1RING
luciferase-osk-3'UTR
luciferase-oskΔAR-3'UTR
2.5% input
IP (FLAG)
α-FLAG
D
FLAG – GFP
FLAG – Mkrn1RING
LacZ KD
Imp KD
luciferase-osk-3'UTR
2.5% input
IP (FLAG)
α-FLAG
Relative gene expression of Imp mRNA
LacZ KD
Imp KD
FLAG - GFP
FLAG - Mkrn1RING
E
FLAG– GFP
FLAG– Mkrn1RING
LacZ KD
pAbp KD
luciferase-osk-3'UTR
2.5% input
IP (FLAG)
α-FLAG
Relative gene expression of pAbp mRNA
LacZ KD
pAbp KD
FLAG - GFP
FLAG - Mkrn1RING
F
FLAG – GFP
FLAG – Mkrn1
FLAG – Mkrn1ΔZnF1
FLAG – Mkrn1PAM2
FLAG – Mkrn1ΔZnF1+PAM2
luciferase-osk-3'UTR
2.5% input
IP (FLAG)
α-FLAG

Dold_Fig_S12

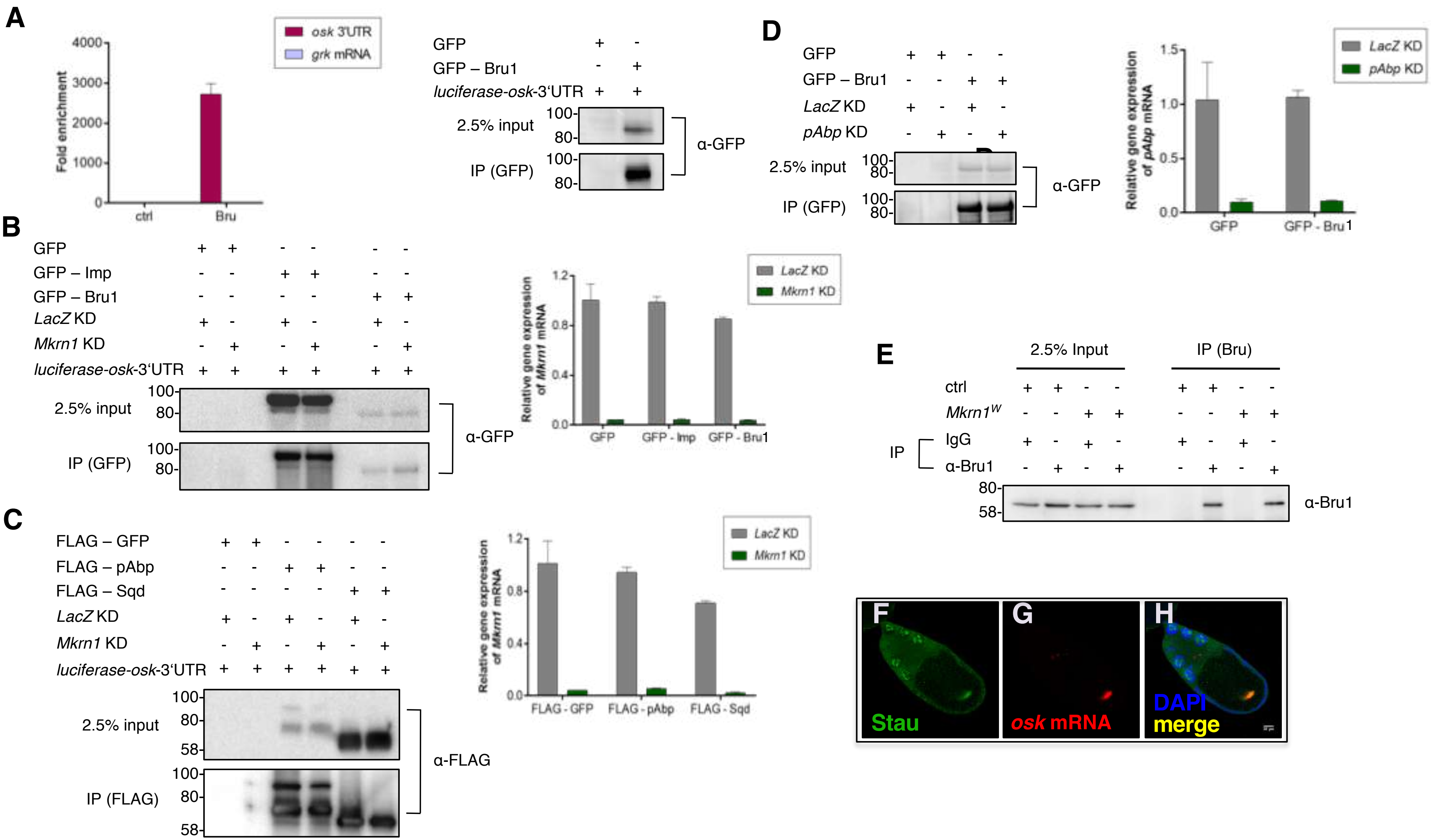
A
osk 3'UTR
grk mRNA
Fold enrichment
4000
3000
2000
1000
0
ctrl
Bru
GFP + -
GFP – Bru1 - +
luciferase-osk-3'UTR + +
2.5% input
IP (GFP)
100-
80-
α-GFP
B
GFP + + - - - -
GFP – Imp - - + + - -
GFP – Bru1 - - - - + +
LacZ KD + - + - + -
Mkrn1 KD - + - + - +
luciferase-osk-3'UTR + + + + + +
2.5% input
IP (GFP)
α-GFP
LacZ KD
Mkrn1 KD
Relative gene expression of Mkrn1 mRNA
1.2
0.8
0.4
0.0
GFP
GFP - Imp
GFP - Bru1
C
FLAG – GFP + + - - - -
FLAG – pAbp - - + + - -
FLAG – Sqd - - - - + +
LacZ KD + - + - + -
Mkrn1 KD - + - + - +
luciferase-osk-3'UTR + + + + + +
2.5% input
IP (FLAG)
100-
80-
58-
α-FLAG
FLAG - GFP
FLAG - pAbp
FLAG - Sqd
D
GFP + + - -
GFP – Bru1 - - + +
LacZ KD + - + -
pAbp KD - + - +
2.5% input
IP (GFP)
α-GFP
pAbp KD
Relative gene expression of pAbp mRNA
1.5
1.0
0.5
0.0
GFP - Bru1
E
2.5% Input
IP (Bru)
ctrl + + - - + + - -
Mkrn1W - - + + - - + +
IP
IgG + - + - + - + -
α-Bru1 - + - + - + - +
80-
58-
α-Bru1
F
G
H
Stau
osk mRNA
DAPI
merge
